# Coordinated phase separation and phase transition underlie synaptic ribbon condensate formation and plasticity

**DOI:** 10.64898/2026.04.28.721253

**Authors:** Yang Liu, Xu Wang, Tianyu Zheng, Fengfeng Niu, Ruiyi Sun, Chao Yang, Shun Xu, Zhipeng Zhao, Zeyu Shen, Weilin Huang, Xuejie Wang, Kai Liu, Shujun Cai, Mingjie Zhang, Zhiyi Wei

## Abstract

Mesoscale molecular apparatuses with diverse functions are widespread in cells. How mesoscale cellular apparatuses are formed and regulated is poorly understood. The synaptic ribbon in sensory neurons, which can dynamically adjust its size and shape in response to light or sound, is a mesoscale molecular assembly. In this study, we demonstrate that RIBEYE, the backbone protein of synaptic ribbons, via combined ordered and disordered interactions, forms micron-sized dynamic molecular assemblies. Integrated cryo-electron microscopy and tomography studies revealed that the N-terminal SAM domain and the C-terminal B domain of RIBEYE form distinct nanoscale filaments and further assemble into ribbon-like architectures via phase transition. Unexpectedly, the N-terminal intrinsically disordered region of RIBEYE softens the solid mesoscale ribbon structures via phase separation. The mesoscale size and shape of the RIBEYE condensate are bidirectionally tuned by physiological modulators such as Piccolino and CtBP1, linking synaptic ribbon structural plasticity to functional adaptability. Thus, our study demonstrates that combined phase transition and phase separation of nanoscale protein oligomers can build mesoscale functional cellular apparatuses.

## Introduction

Living cells contain numerous mesoscale subcellular structures or molecular apparatuses, usually with micrometers in their sizes. Such mesoscale molecular apparatuses dynamically adjust their morphologies to exert their functions in response to stimuli. Among these, membrane-demarcated organelles, such as the endoplasmic reticulum and Golgi apparatus, have been extensively studied and are relatively well understood ^1–4^. Another distinct class of subcellular structures found in cells is not membrane-enclosed but instead formed by autonomous assembly of biomolecules into condensed assemblies (referred to as membraneless apparatuses for simplicity). Such mesoscale membraneless apparatuses include synaptic ribbons in sensory neurons ^5–8^, axon initial segments at the beginning of axons of neurons ^9,10^, focal adhesions and podosomes in migrating cells ^11–14^, costameres, Z-Disc, and dense bodies in muscles ^15–17^, *etc*. The molecular mechanisms and structural bases underlying the autonomous assembly and dynamic regulation of such membraneless apparatuses are poorly understood, as the sizes of these apparatuses fall into a window that is not easily accessible by currently available techniques. Nonetheless, liquid-liquid phase separation (LLPS) and liquid-to-solid phase transition have been implicated in forming some of these mesoscale structures ^18–21^.

Synaptic ribbon, identified over six decades ago ^22^, is a special type of presynaptic active zone (AZ) found in synapses of sensory neurons such as retinal photoreceptors and inner ear hair cells ^8,23–25^. Under optical or electron microscopes, the AZ of retinal photoreceptors appeared as a micron-sized, horseshoe/plate-like ribbon ^22,25^, so the name of synaptic ribbon was given (see Fig. 1a for a scheme). The morphological plasticity is a fascinating feature of synaptic ribbons. For instance, in nocturnal rodents like mice and rats, retinal synaptic ribbons are known to undergo light-dependent structural remodeling. In a dark environment, such as in the evening, synaptic ribbons in rodent photoreceptors expand their physical sizes (Fig. 1a) ^26–28^. This enlarged morphology allows more synaptic vesicles to dock onto the AZ for increased and sustained neurotransmitter release ^29^, thereby improving the animal’s vision under dim light. Conversely, exposure to daylight induces structural changes of the ribbons marked by the formation of droplet-like AZ fragments at the ribbon apex, resulting in shrinkage of membrane-attached synaptic ribbons and consequent reduction in photoreceptor sensitivity (Fig. 1a) ^26–28^. Synaptic ribbons of photoreceptors of hibernating animals, such as ground squirrels, almost completely disappear during hibernation as their photoreceptors are essentially switched off ^30,31^. The mechanistic basis governing such light-dependent mesoscale remodeling of synaptic ribbons is not understood.

**Figure 1.**
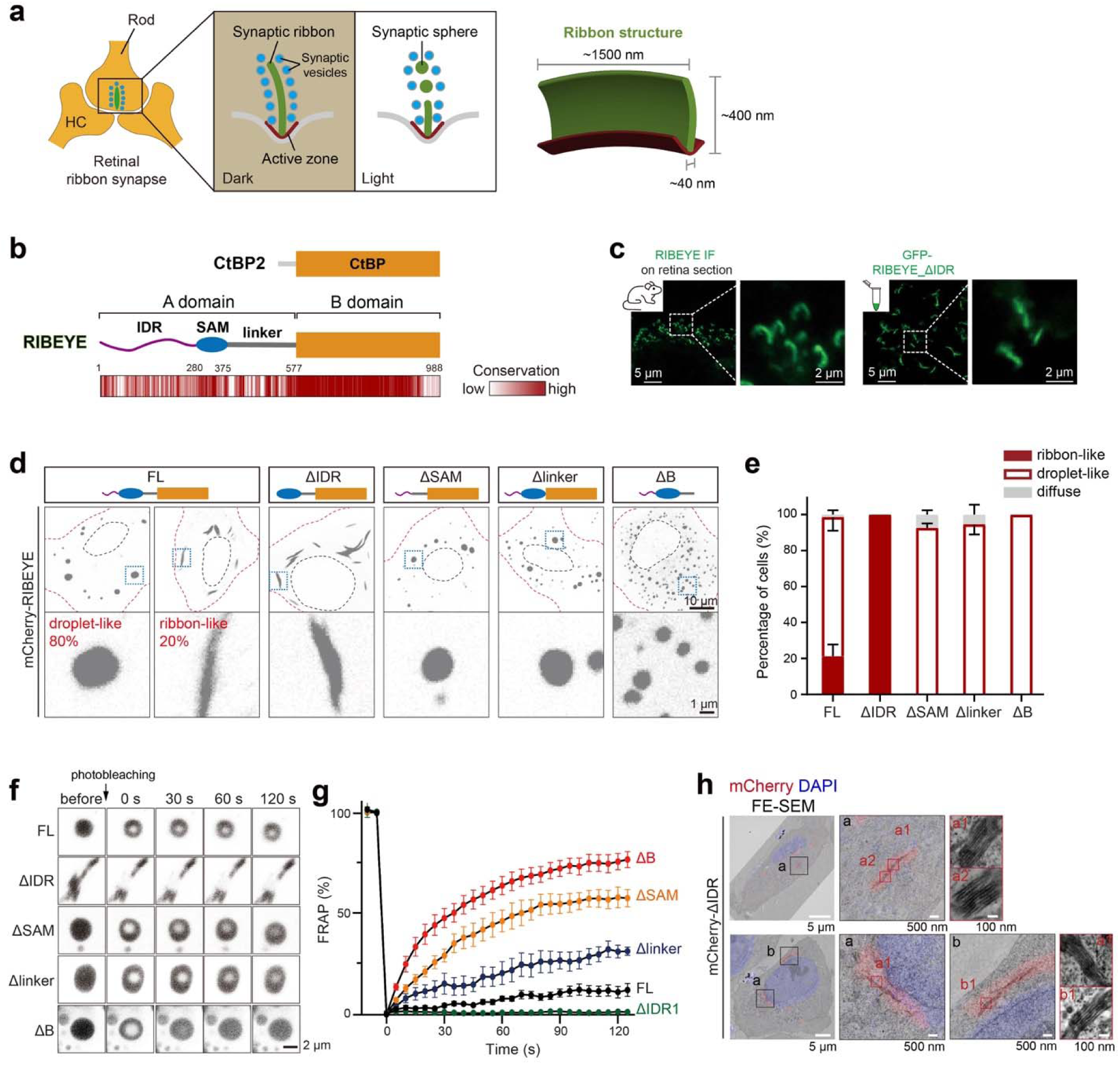
RIBEYE can form both solid ribbon-like and liquid-like condensates. **a** Schematic of a ribbon synapse in photoreceptors. The synaptic ribbon (green) is attached to the AZ (red) at the presynaptic terminal. Synaptic vesicles (blue) surround the ribbon structure. The structure of the synaptic ribbon is regulated by light conditions. In the dark, this structure adopts a ribbon-like shape; in a light environment, the structure undergoes a morphological transformation into a spherical shape. A cartoon representation of the ribbon structure in a rod photoreceptor is shown in the right panel. **b** Domain organization of CtBP2 and its splicing isoform RIBEYE. The sequence conservation of RIBEYE was generated through a multi-sequence alignment of RIBEYE sequences from various vertebrates, including human, mouse, bovine, panda, whale, alligator, snake, frog, zebrafish, lamprey, coelacanth, and salmon. **c** The formation of ribbon-like structures by RIBEYE *in vivo* and *in vitro*. The protein concentration of GFP-RIBEYE_ΔIDR is 10 μM. **d** HeLa cell imaging of droplet-like or ribbon-like condensates formed by different RIBEYE constructs. Representative condensates are shown as enlarged views below the corresponding transfected cells. **e** Quantification of the percentage of transfected cells showing different condensates in panel **d**. n = 3 repeats, in each repeat 20-40 cells are quantified. The estimate of variation is indicated by the s.d.. **f** Representative images of FRAP experiments with RIBEYE and its truncations in HEK293T cells. **g** FRAP analysis of condensates formed by RIBEYE and its truncations. n = 5-10 cells for each curve. The estimate of variation is indicated by the s.e.m.. **h** FE-SEM imaging of HeLa cells transfected with RIBEYE_ΔIDR showing filamentous structures in ribbon-like condensates.

The formation of synaptic ribbons depends on RIBEYE, which is specifically expressed in neurons forming ribbon synapses ^32–34^. RIBEYE was identified as a splicing variant of a nuclear transcriptional co-repressor, termed C-terminal binding protein 2 (CtBP2) ^34^. Structurally, RIBEYE contains an evolutionarily divergent A domain and a conserved B domain, with the latter being identical to the CtBP2 core domain (Fig. 1b) ^34^. Previous biochemical and structural investigations have demonstrated that the core domain mediates CtBP2 tetramerization ^35,36^, suggesting that the B domain plays a role in RIBEYE-mediated synaptic ribbon assembly ^37^. However, recent studies have suggested that RIBEYE may drive the synaptic ribbon formation through homomeric interactions involving its A domain ^38,39^, though how the A domain assembles into homotypic multimers and how this multimerization process contributes to the formation of the mesoscale synaptic ribbon structure remain unclear.

In this study, we dissected the dynamic assembly of RIBEYE and uncovered its dual ability to form both ribbon-like structures and droplet-like condensates. Cryo-electron microscopy (cryo-EM) reveals that the A and B domains of RIBEYE each contain conserved structural motifs enabling the self-assembly of distinct helical filaments. Cryo-electron tomography (cryo-ET) further elucidated how these nanoscale filaments, as basic units, interlock to generate a mesoscale solid architecture of the synaptic ribbon. Concurrently, the N-terminal intrinsically disordered region (IDR) in the A domain drives LLPS of this basic unit, providing the mesoscale synaptic ribbon with flexibility and fluidity. Such mesoscale synaptic structures are dynamic and bidirectionally modulated by physiological regulators, Piccolino and CtBP1, through tuning the phase separation or phase transition properties of the RIBEYE condensate. The findings on the formation and plastic regulation of the synaptic ribbon may serve as a model for understanding other cellular mesoscale apparatuses self-assembled via molecular condensation.

## Results

### RIBEYE is capable of assembling into both ribbon and spherical-shaped condensates

Although the A domain of RIBEYE lacks canonical structural motifs as suggested by sequence analysis, an AlphaFold-based structure prediction reveals a sterile alpha motif (SAM) domain in the middle of the A domain, flanked by an N-terminal IDR and a C-terminal flexible linker connecting the SAM domain and the B domain (Fig. 1b; Supplementary Fig.S1a). Guided by this predicted model, we removed the N-terminal IDR region and successfully purified the RIBEYE_ΔIDR protein with an MBP-GFP tag (Supplementary Fig. S1b). Strikingly, following MBP cleavage, purified RIBEYE_ΔIDR spontaneously assembled into ribbon-like structures, resembling the ultrastructural shape of native synaptic ribbons stained by anti-RIBEYE antibody in murine photoreceptors (Fig. 1c). When transiently expressed in mammalian cells lacking endogenous RIBEYE, mCherry-tagged full-length protein (RIBEYE_FL) predominantly formed spherical condensates as previously reported ^34,40^, but with a significant portion forming ribbon-like structures (Fig. 1d, e; Supplementary FigS1c). In contrast, the ΔIDR mutant exclusively generated elongated, solid-like condensates, morphologically like synaptic ribbons (Fig. 1d, e; Supplementary Fig. S1c). RIBEYE mutants with deletions of other individual structural elements (e.g., the SAM domain, the linker, or the B domain), instead, formed droplet-like condensate (Fig. 1d, e; Supplementary Fig. S1c). Each of these isolated domains or structural segments exhibited cytoplasmic dispersions in heterologous cells (Supplementary Fig. S1d).

Next, we investigated the material properties of the RIBEYE condensates. Live cell imaging captured dynamic fusion events in spherical droplets formed by RIBEYE_FL, though with relatively slow fusion speed (Supplementary Fig. S1e). In contrast, the ribbon-like structures formed by RIBEYE_ΔIDR were never observed to coalesce, suggesting that such ribbon-like structures observed in test tubes and in heterologous cells are solid-like structures formed by the phase transition of RIBEYE_ΔIDR. Fluorescence recovery after photobleaching (FRAP) analysis further showed complete immobility of the RIBEYE_ΔIDR condensate, indicative of a solid-like phase of the condensate (Fig. 1f, g; Supplementary Fig. S1f). The solid assemblies formed by RIBEYE_ΔIDR are not irreversible protein aggregates, as these assemblies can be dynamically modulated by physiological regulators such as Piccolino and CtBP1 (see later part of the manuscript). Curiously, molecules in the RIBEYE_FL condensate have limited molecular mobility (∼9% mobile fraction) even though the condensate exhibits spherical shape and macroscopic fluidity, evidenced by droplet fusions (Supplementary Fig. S1e). Interestingly, the deletion of the SAM or B domain dramatically increases the mobile fraction of condensates to over 60%. Deleting the entire linker connecting the SAM domain and the B domain also increased condensate dynamics (Fig. 1f, g; Supplementary Fig. S1f). Field emission scanning electron microscopy (FE-SEM) on chemically fixed cells expressing RIBEYE_ΔIDR revealed highly organized RIBEYE filaments assembled into ribbon-shaped supramolecular structures through lateral stacking (Fig. 1h). Taken together, the above biochemical and biophysical studies revealed that RIBEYE is capable of forming mesoscale filament-like structures resembling synaptic ribbons via phase separation and phase transition. Every structural element of the protein contributes to the morphology and material property of the formed RIBEYE condensate.

### The SAM and B domains of RIBEYE assemble into helical filaments as the basic units of synaptic ribbons

To determine the self-assembly capability of the predicted SAM domain in RIBEYE, we purified a RIBEYE fragment containing the SAM domain. Analytical size-exclusion chromatography indicated that the SUMO-tagged SAM fragment exists primarily as monodisperse monomers (Supplementary Fig. S2a). Upon tag cleavage, we found that the SAM fragment forms reversible precipitates (Supplementary Fig. S2b), which were confirmed as filaments by negative-stain EM (nsEM) (Fig. 2a), indicating the SAM domain can undergo phase transition from homogeneous dilute solution to solid filaments. Cryo-EM structural determination of the SAM filaments at ∼3.4 Å resolution using single particle analysis (SPA) method (Supplementary Fig. S3 and Table S1) revealed two double-helical architectures: a major 6 nm-diameter filament (Fig. 2b; thick-form at ∼90%) and a minor 4.5 nm-diameter variant (Fig. 2c; thin-form at ∼10%), both exhibiting a -36° helical twist with 10 SAM domain pairs per turn. These helical architectures use similar dual SAM/SAM interaction modes for the filament assembly (Movie S1). First, SAM fragments stack along the helical axis via the canonical head- to-tail SAM/SAM interaction (Supplementary Fig. S2c), involving both charge-charge (e.g., salt bridges formed by E294 and R334) and hydrophobic interactions (e.g., F309, V314, T317, L330, and V335). Second, two helical SAM protofilaments associate laterally through the extended C-terminal α-helix (α5) of the SAM domain, which is uniquely long in RIBEYE SAM (Fig. 2d, a magnified view). Structural comparison between the thick and thin forms showed divergence in the packing angle between the paired α5-helices (Supplementary Fig. S2d). It is likely that the weak association between α5-helices enables angle changes, allowing the dynamic transition between the two SAM filament forms (Movie S2). In alignment with the above structural analysis, mutations of selected amino acid residues designed to interfere with the SAM/SAM interactions severely disrupted filament assembly as evident by nsEM analysis (Supplementary Fig. S2e).

**Figure 2.**
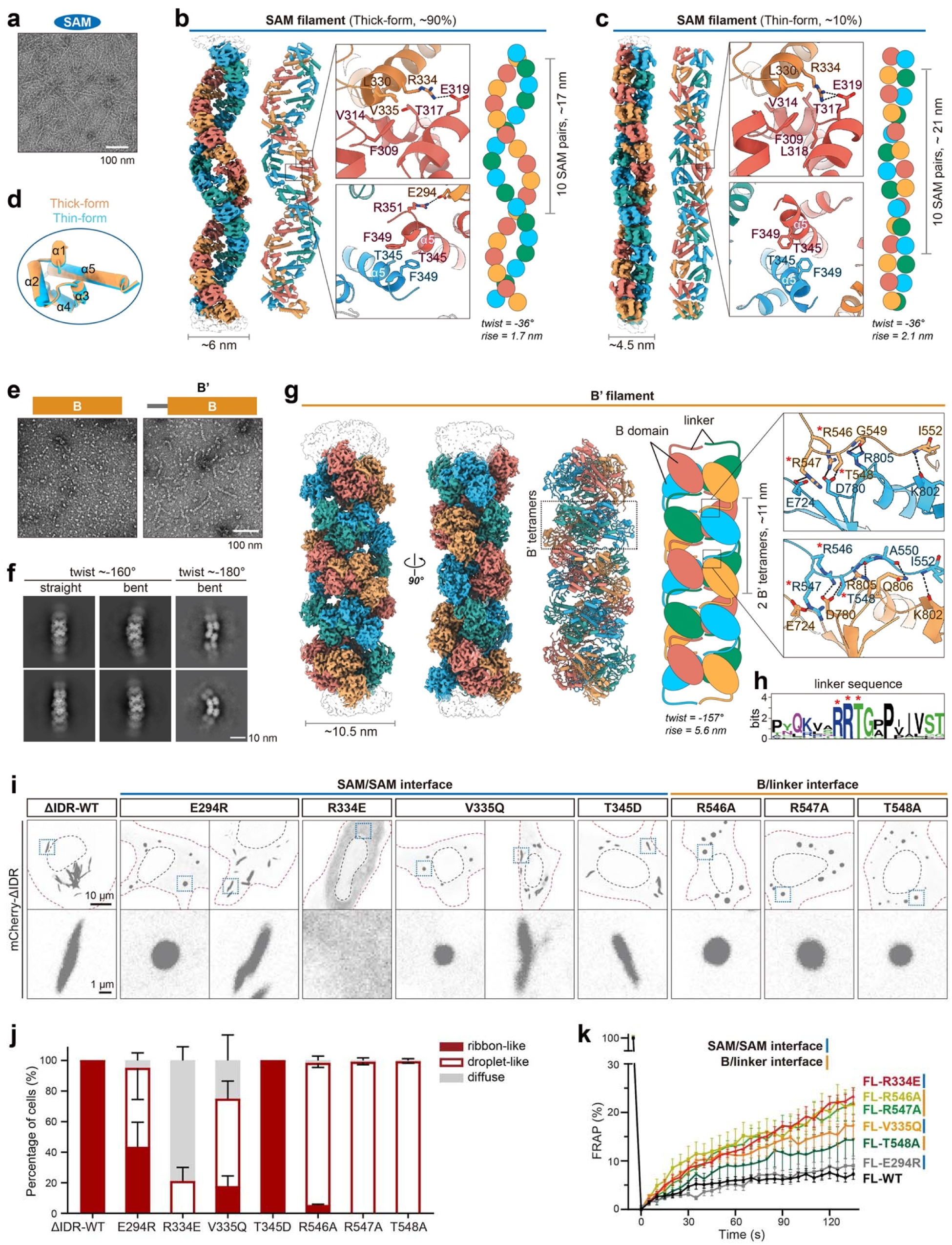
Structural characterization of two filament assemblies in RIBEYE. **a** NsEM analysis showing the filament structures formed by purified RIBEYE SAM fragments. **b-d** Cryo-EM structures of the SAM filament. The density maps, atomic models, two SAM/SAM interfaces for double helical packing, and cartoon models of the SAM filament in both thick (**b**) and thin (**c**) forms are presented from left to right. Both filaments are assembled from SAM protomers adopting an identical conformation (**d**). **e** NsEM analysis of purified B and B’ fragments. The addition of the N-terminal linker region in B’ fragments induces filament formation. **f** Cryo-EM analysis of the B’ filament. The representative 2D classes of straight and bent filaments with different twist angles were displayed. **g** Cryo-EM structure of the B’ filament. The density map, atomic model, cartoon model, and two similar linker/B interfaces of the B’ filament are presented from left to right. The three key interface residues, R546, R547, and T548, in the linker region are labeled by red stars. **h** Sequence analysis showing the highly conserved feature of the RRT motif in the linker sequences from different species. The plot was generated using WebLogo (https://weblogo.berkeley.edu). **i** HeLa cell imaging of condensates formed by RIBEYE_ΔIDR with either SAM/SAM or B/linker interface mutations. **j** Quantification of condensate morphology in cells shown in panel **h**. n = 3 repeats, in each repeat 20-40 cells are quantified. The estimate of variation is indicated by the s.d.. **k** FRAP analysis of condensates formed by RIBEYE_ΔIDR with mutations at either SAM/SAM or B/linker interfaces. n= 5-10 cells for each curve. The estimate of variation is indicated by the s.e.m..

As the A domain and B domain of the RIBEYE were implicated to interact with each other ^38^, we sought to map the B-interacting region in the A domain. Our investigation revealed that, although no direct interaction between the SAM and B domains could be detected, a short stretch of sequence immediately preceding the N-terminus of the B domain specifically binds to the B domain with a *K*_d_ of ∼7 μM (Supplementary Fig. S2f). Strikingly, extending the B domain boundary to include this link sequence (referred to as B’ hereafter) converts the globular B domain tetramer into filamentous B’ oligomers with substantial conformational flexibility (Fig. 2e, f; and Supplementary Fig. S2g). We determined the straight form of the B’ filament structure at 3.41 Å resolution (Supplementary Fig. S4 and Table S1). In this filament structure, the main core corresponding to the B domain boundary forms tetramers in a manner identical to the CtBP proteins (Supplementary Fig. S2h). These B’ tetramers stack on top of each other via their short axis (defined as the vertical side) by inter-tetramer B domain/linker interaction to form elongated B’ filaments (Fig. 2g and Movie S3). Specifically, each B’ tetramer interacts with two linkers from an upper B’ tetramer and two linkers from a lower B’ tetramer (Fig. 2g, right cartoon panel; Supplementary Fig. S2i). Interestingly, the B/linker interaction does not involve the well-characterized hydrophobic cleft at the lateral side of the tetramer that mediates the B domain’s binding to targets with a pentapeptide motif containing the “PXDLS” sequence, where ‘X’ denotes any common amino acids (Supplementary Fig. S2h) ^41,42^. Instead, the B’ filament formation is mediated by a distinct region of the B domain, which recognizes a highly conserved RRT motif (R546, R547, and T548) in the linker sequence by forming salt bridges and hydrogen bonds (Fig. 2g, h, magnified inserts). Mutating any one of these three residues in this motif to alanine abolished the B’ filament formation (Supplementary Fig. S2j). Interestingly, this binding mode was previously indicated in the binding of CtBP1 to transcription factors that contain similar RRT motifs (Supplementary Fig. S2h) ^43^, suggesting a conserved RRT-binding mode by the B domain.

To confirm the role of the SAM and B’ filaments in RIBEYE assemblies, we introduced the filament-disrupting mutations targeting the SAM/SAM or B/linker interface into RIBEYE_ΔIDR and expressed these mutants in heterologous cells. Confocal imaging analysis revealed that most mutations impaired ribbon-like condensate formation and converted RIBEYE_ΔIDR into droplet-like condensates or even a largely diffused distribution for the R334E mutant (Fig. 2i, j), suggesting that both the SAM and B’ filament formations are required for the ribbon-like structure assembly of RIBEYE_ΔIDR. The only exception was the T345D mutant, which compromises the lateral SAM/SAM interaction via the α5-helix packing (Fig. 2b) and thus likely converts the SAM domain into a single-helix filament instead of the double-helix filament. This result suggests that the filamentous oligomerization of the SAM domain, but not specifically the double-helix filaments, is essential for the RIBEYE_ΔIDR ribbon structure formation. In addition to the formation of ribbon-like structures, the filamentous assemblies of the SAM and B’ domains also contribute to the low dynamic property of droplet-like RIBEYE condensates, as RIBEYE variants harboring mutations at the SAM/SAM or B/linker interfaces showed increased molecular mobility in condensates formed by the mutants, assayed by FRAP experiments (Fig. 2k; Supplementary Fig. S2k).

### The mesoscale ribbon-like structure of RIBEYE is formed via a high-order assembly of the SAM and B’ filaments

To investigate how SAM and B’ filaments assemble into ribbon-like structures, we performed cryo-ET of GFP-tagged RIBEYE_ΔIDR condensates in cells, using GFP fluorescence signal as the guide for positioning ribbon-like structures during cryo-focused ion beam (cryo-FIB) milling (Supplementary Fig. S5a-c). Cryotomograms revealed ∼10 nm thick filaments, extending over 800 nm in length, which are packed side-by-side into flat, sheet-like assemblies (Fig. 3a, b; Supplementary Fig. S5d). We defined three orthogonal views of these assemblies as front, side, and top views (Fig. 3b and Movie S4). Multiple such sheets are stacked in parallel, with an inter-sheet spacing of ∼40 nm (Fig. 3b; Supplementary Fig. S6a), measured from the centers of the filaments (this convention is consistently applied throughout subsequent measurements). These observations indicate an ordered, layered architecture within the ribbon-like condensates formed by RIBEYE_ΔIDR. Subtomogram averaging (STA) of individual filament segments was achieved *in situ* at 16.9 Å resolution (Fig. 3c; Supplementary Fig. S7a). Although the cryo-EM structure of the straight B’ filament shares a similar diameter with the STA density map (∼10 nm), it showed poor map fitting (Supplementary Fig. S7b). Helical symmetry analysis of the STA map suggested a ∼-179° twist and ∼6.3 nm rise (Supplementary Fig. S7a), a symmetry resembling that observed in some B’ filaments assembled from purified B’ fragments (Fig. 2f; Supplementary S2g). By incorporating this symmetry, the refined B’ filament model provides a robust fit to the STA density map (Fig. 3c; Supplementary Fig. S7c and Movie S5). We thus refer to these layered assemblies as the B’ sheets, in which inter-filament interactions may stabilize such a B’ helical arrangement.

**Figure 3.**
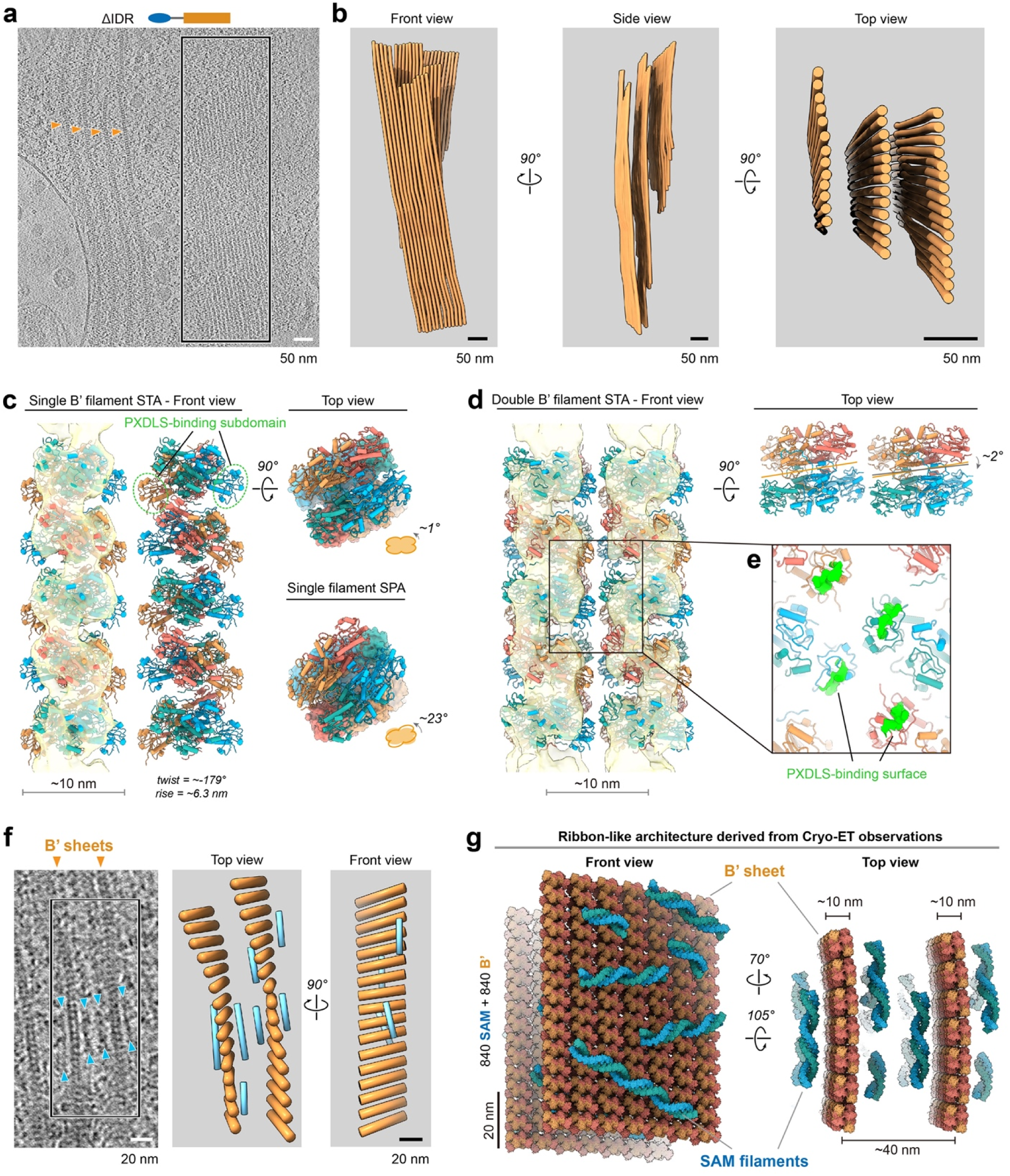
*In situ* reconstitution of the ribbon-like architecture. **a** Cryotomographic slice of a HeLa cell exogenously expressing GFP-tagged RIBEYE_ΔIDR. The two different orientations of sheet-like assemblies of RIBEYE_ΔIDR were indicated by orange arrowheads (top view) and boxed (front view), respectively. **b** Front, side, and top views of 3D renderings of sheet-like assemblies boxed in panel **a**. **c** STA density map and model of a single B’ filament. The map is shown fitted with the B’ filament model assembled from B’ tetramers with indicated helical twist and rise. The twist angles between the B’ filament models determined by STA and SPA are compared. **d** STA density map and overlaid model of two adjacent B’ filaments. This double filament model was generated by rigid-body fitting the single filament shown in panel **c** into the density map. The distance and rotation angle between the two filaments are indicated. **e** A zoomed-in view of PXDLS-binding subdomains in B domains that mediate inter-filament packing. Upon this packing, the two PXDLS-binding surfaces are still available for target binding. **f** Cryotomographic view of perpendicularly oriented B’ sheets. This represents a top view of two neighboring B’ sheets in panel **b**. The B’ and SAM-like filaments are indicated by orange and cyan arrowheads in the left panel, respectively. The corresponding 3D rendering of B’ filaments (orange) and SAM-like filaments (cyan) is presented in the right panel. **g** An atomic model showing the ribbon-like architecture assembled by SAM and B’ filaments of RIBEYE. The model, generated based on the cryo-ET observations, contains a total of 840 SAM domains and 840 B’ domains. The thick-form SAM filaments, dominant in our cryo-EM analysis, were used as the SAM filament component in this model. The flexible regions in RIBEYE, including the IDR and Linker, are excluded from the model.

To understand how B’ filaments are bundled regularly into sheets, we performed STA on pairs of adjacent filaments, focusing on inter-filament interactions (Supplementary Fig. S7d). The resulting STA map showed a side-by-side arrangement of B’ filaments, with the distance between neighboring B’ filaments measured at ∼10 nm (Fig. 3d; Supplementary Fig. S6b). By fitting the B’ filament obtained from Fig. 3c into the map, we generated a double filament model to elucidate the inter-filament interaction (Fig. 3d and Movie S6). In this model, the two filaments are oriented in parallel, with a subtle rotation along their helical axis (Fig. 3d). Given the 2-fold symmetry in the B tetramer, a near 180° twist observed in the STA-fitted B’ filament aligns the PXDLS-binding subdomains on the two sides of the filament (Fig. 3c). This specific spatial arrangement in the double filament model allows a consistent packing interaction, mediated by two neighboring PXDLS-binding subdomains (Fig. 3d), for the organization of the entire B’ sheet. Notably, the PXDLS-binding sites are completely accessible in this architecture (Fig. 3e), suggesting that the binding of PXDLS motifs could still occur upon B’ sheet formation.

Interestingly, in a region adjacent to the front-view area, we observed B’ sheets oriented perpendicularly to the front views (i.e., arrowhead highlighted objects vs the boxed region in Fig. 3a). Such perpendicularly oriented B’ sheets likely represent the B’ sheets in the top view (Fig. 3f and Movie S7). A low-magnification projection capturing these top views suggests that B’ sheets can extend to at least 3.5 µm in width (Supplementary Fig. S5d). Given the inter-B’ filament distance of ∼10 nm, we estimated that each B’ sheet may consist of over 350 B’ filaments packed side-by-side, suggesting a high degree of stability in forming ribbon-like structures via higher-order packing of B’ sheets in the RIBEYE_ΔIDR condensate.

Compared to the curved, short filaments (∼60-100 nm in length, Fig. 2e, f) formed by the purified B’ domain, the B’ filaments in ribbon-like condensates formed by RIBEYE_ΔIDR show enhanced stability and extended length, suggesting that the SAM domain can facilitate the assembly of the short B’ filaments into stable and mesoscale ribbon-like structures. Interestingly, we observed the presence of short, discontinuous filamentous densities with ∼6 nm in thickness, on the two sides of each B’ sheet (Fig. 3f). These filaments were oriented nearly perpendicular to the B’ filaments (Fig. 3f). The diameter of these short filaments matches well with that of the major SAM filament resolved by cryo-EM (Fig. 2b). Thus, we propose that these filaments are likely SAM filaments, although their low abundance and high flexibility precluded meaningful STA analysis. Notably, the morphology of layered B’ sheets flanked by these SAM-like filaments closely resembles the EM images of synaptic ribbons reported previously ^44^. Our cryo-ET analysis determined that two neighboring B’ sheets are regularly spaced by ∼40 nm (Fig. 3f; Supplementary Fig. S6a), which is close to the thickness of each synaptic ribbon from rat retinal synapses measured by EM ^40^. It is likely that synaptic ribbons form from two proximately apposed B’ sheets, spaced by SAM filaments that may link the two B’ sheets together. Together, our *in situ* cryo-ET analysis of RIBEYE_ΔIDR offers an atomic view showing the unique 3D organization of the B’ and SAM domains into mesoscale synaptic ribbon-like structures (Fig. 3g and Movie S8).

### The N-terminal IDR enhances the liquidity of the RIBEYE condensate by suppressing filament growth

We noticed with interest that the N-terminal IDR in the full-length RIBEYE can suppress long and bundled filament formation by shifting the RIBEYE condensate towards more fluid and droplet-like morphology (Fig. 1d). Indeed, amino acid sequence analysis of the IDR revealed that the region has a high predicted phase separation score (Supplementary Fig. S8a), suggesting that the IDR has a LLPS capacity. Although the IDR alone does not form cellular condensates (Supplementary Fig. S1d), fusion of the IDR with the SAM domain or with the B domain can massively promote phase separation of each of the two domains (Supplementary Fig. S8b). Consistently, *in vitro* biochemical experiments demonstrated that the purified IDR displayed a concentration-dependent phase separation in the presence of a low concentration of a crowding reagent (3% PEG-8000) (Fig. 4a), and the IDR condensate is soft and dynamic (Supplementary Fig. S8c, d). We further narrowed down the N-terminal half of the IDR to be essential for forming droplet-like RIBEYE condensates, as deleting this region is sufficient to shift condensate morphology toward ribbon-like structures (Supplementary Fig. S8e, f). Sequence analysis of this region uncovered 14 conserved aromatic residues (Supplementary Fig. S8g), suggesting π-π interactions as a key determinant of the IDR-mediated LLPS. To validate this, we systematically mutated varied numbers (3-14) of these aromatic residues to alanine and found that as the number of aromatic residues substituted, the LLPS capability of the purified IDR decreased. No liquid droplets could be observed in the 11A and 14A IDR mutants (Fig. 4b). Mirroring these biochemical findings, RIBEYE_FL bearing these mutations exhibited a progressive conversion from liquid-like to ribbon-like condensates in cells when increasing numbers of aromatic residues in the IDR were substituted by alanine (Fig. 4c, d). Thus, we concluded that aromatic-mediated multivalency in the IDR contributes to maintaining RIBEYE condensate fluidity.

**Figure 4.**
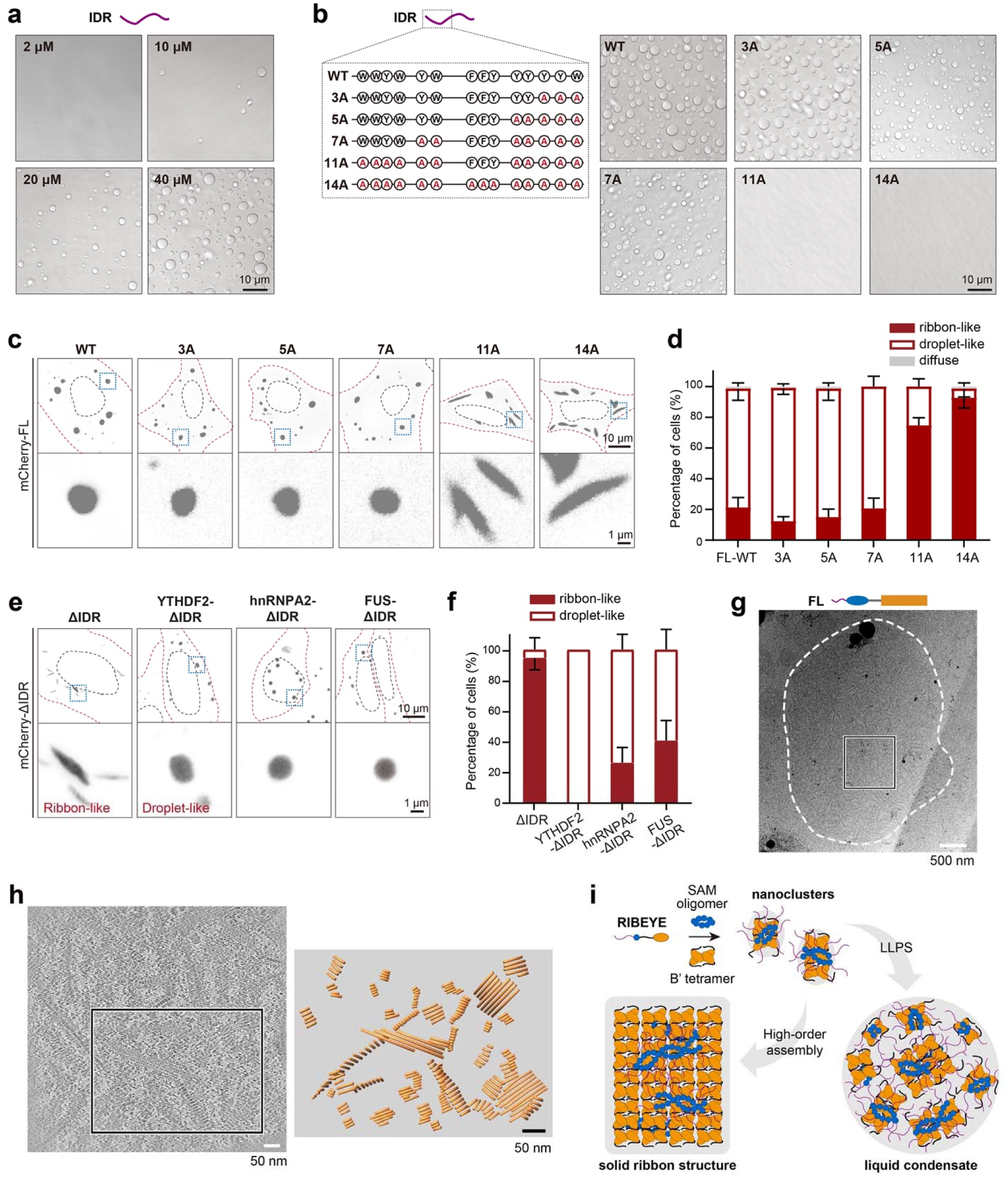
RIBEYE’s IDR promotes LLPS and regulates the RIBEYE phase transition. **a** Representative images of RIBEYE_IDR droplet formation at different concentrations in the presence of 3% (w/v) PEG 8000. **b** Representative images of RIBEYE_IDR (40 μM) droplet formation with aromatic amino acid to alanine mutations in the IDR. The design of these mutations is schematically illustrated. A crowding reagent of 3% (w/v) PEG 8000 was applied. **c** HeLa cell imaging of condensates formed by RIBEYE with different alanine mutations in the IDR. **d** Quantification of condensate morphology in cells shown in panel **c**. n = 3 repeats, in each repeat 20-40 cells are quantified. The estimate of variation is indicated by the s.d.. **e** HEK293T cell imaging of condensates formed by chimeric RIBEYE constructs with their IDR sequence replaced by other pro-LLPS sequences. **f** Quantification of condensate morphology in cells shown in panel **e**. =3 repeats, in each repeat 30-50 cells are quantified. The estimate of variation is indicated by the standard error of the s.e.m.. **g** Low-magnification cryo-EM projection of a cryolamella (see Supplementary Fig. S9) from a HEK293T cell exogenously expressing mCherry-tagged RIBEYE_FL. **h** Zoomed-in cryotomographic view of the boxed region in panel **g**. The corresponding 3D rendering of disorganized B’ sheets is presented in the right panel. **i** A cartoon model illustrating the RIBEYE nanocluster-mediated phase separation and transition. These RIBEYE nanoclusters are preassembled via oligomerization of the SAM and B domains, forming short filaments. These nanoclusters may undergo two different assembly modes: they either transition into solid ribbon structures through regular inter-filament interactions, or form liquid condensates via IDR-mediated LLPS.

To further probe the specificity of IDR-mediated LLPS in counteracting filament-driven ribbon assembly, we engineered chimeric RIBEYE variants by replacing their native IDRs with well-characterized LLPS-promoting IDRs from YTH N6-methyladenosine RNA binding protein F2 (YTHDF2), heterogeneous nuclear ribonucleoprotein A2 (hnRNPA2), or fused in sarcoma (FUS). Despite being physiologically unrelated to RIBEYE IDR, all chimeric constructs tested induced a dramatic increase in droplet-like condensate formation compared to the ΔIDR mutant in cells (Fig. 4e, f). However, a significant portion of cells expressing these chimeras retained ribbon-like condensates (Fig. 4f), suggesting that the native RIBEYE IDR contains evolutionarily refined sequences that optimize LLPS efficiency beyond generic disordered sequences.

The above analysis reveals that the phase separation capacity of the IDR can modulate the mesoscale condensate assembly mode of RIBEYE. Specifically, the IDR may actively suppress uncontrolled growth of mesoscale bundled RIBEYE filaments by promoting phase separation to redirect assembly toward droplet-like condensates. It should be noted that the IDR- mediated suppression of the RIBEYE phase transition toward solid, ribbon-like condensates is not merely a result of the IDR acting as a passive spacer, as the 14A mutant of RIBEYE is incapable of forming the droplet-like condensate (Fig. 4c, d). We propose that the IDR provides weak, heterogeneous, and non-directional multivalent interactions that antagonize the directional growth of the SAM and B’ filaments. To validate this hypothesis, we performed cryo-ET analysis on droplet-like condensates formed by RIBEYE_FL (Supplementary Fig. S9a-c). Our result revealed the presence of small, distorted B’ sheets irregularly distributed within some of these condensates (Fig. 4g, h and Movie S9), suggesting a coexistence of ordered and disordered arrangements of RIBEYE molecules. Furthermore, disrupting filament formation, through deletion of the SAM domain, the linker, or the B domain (Fig. 1g; Supplementary Fig. S1f), or mutation of residues critical for filament formation (Fig. 2k; Supplementary Fig. S2k), enhances droplet liquidity as indicated by increased molecular mobility. Hence, in RIBEYE_FL, these IDR-mediated interactions limit the unregulated high-order assembly of nanoscale filamentous oligomers of the SAM and B’ domains, but instead promote these nanoscale filamentous oligomers into anisotropic and mesoscale condensates (see Fig. 4i for a model).

### Piccolino promotes RIBEYE phase transition and synaptic ribbon formation via stabilizing high-order B’ filament assembly

We next asked how the mesoscale structure of synaptic ribbons could be dynamically regulated to adapt to different signal inputs. Emerging evidence indicates that Piccolino, a ribbon-synapse-specific isoform of the AZ protein Piccolo and a key component of synaptic ribbons ^45–47^, may interact with RIBEYE to modulate the ribbon structure ^40^. By dissecting retinal sections from mice at different developmental stages (postnatal days 14, 21, and 35), we found a positive correlation between the maturation of ribbon synapses and the targeting of Piccolino to the RIBEYE condensates (Fig. 5a, b). Interestingly, this increase in Piccolino levels in RIBEYE condensates was accompanied by a concurrent morphological transition of RIBEYE condensates from droplet-like condensates to the characteristic ribbon shape, a hallmark of mature ribbon synapses (Fig. 5a; Supplementary Fig. S10a), suggesting that Piccolino plays a role in promoting the formation and maturation of synaptic ribbons.

**Figure 5.**
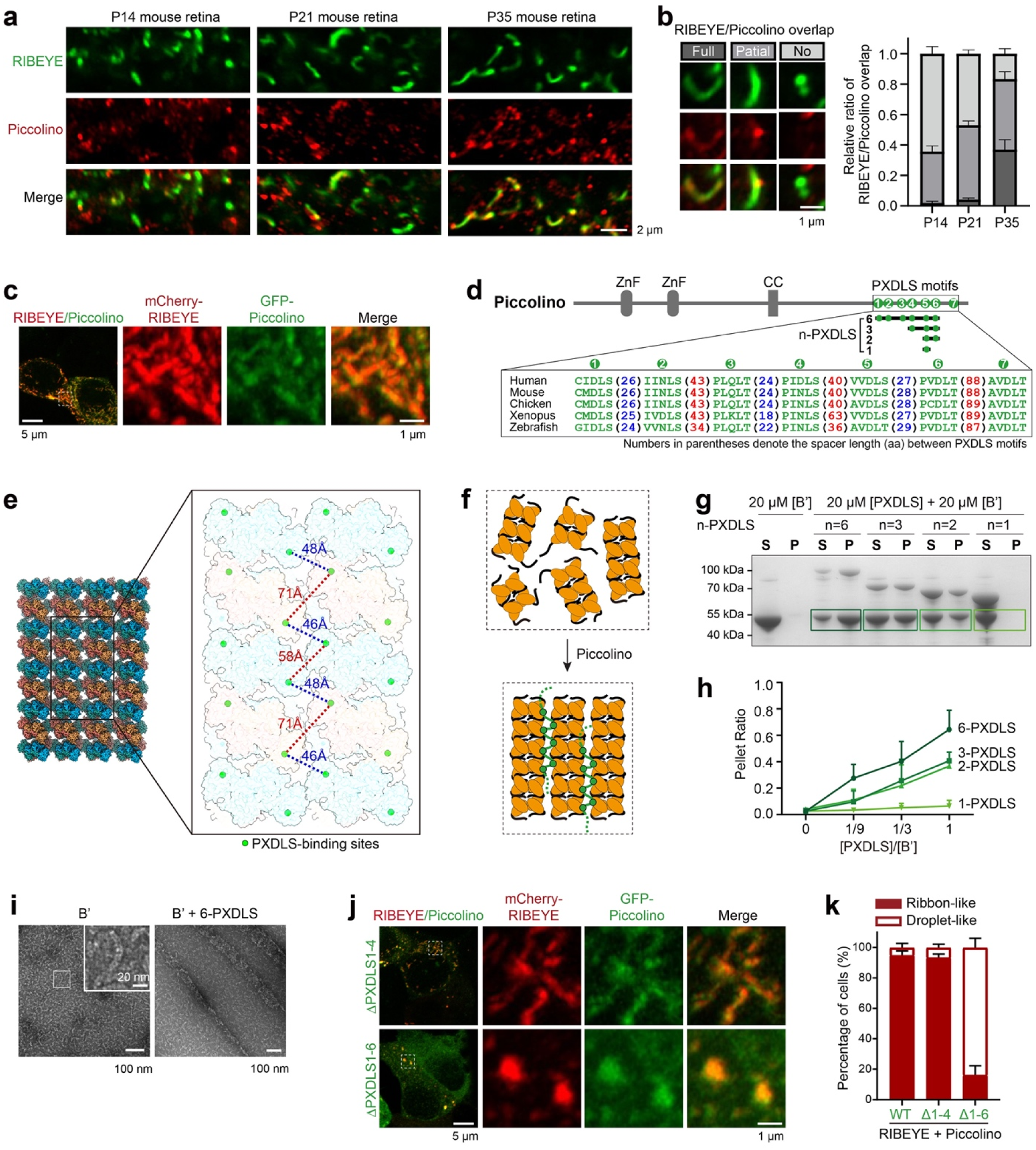
Piccolino promotes ribbon-like condensate formation via its multiple PXDLS motifs. **a** Representative images of the retinal OPL from mice in respective developmental stages (P14, P21, and P35). Sections were stained with Piccolino (red) and RIBEYE (green) antibodies. **b** Quantification of the Piccolino-RIBEYE colocalization in panel **a**. n = 4 animals. The estimate of variation is indicated by the s.e.m.. Zoomed-in views on the left panel show the criteria for the quantification of the Piccolino colocalization with RIBEYE. **c** Cell imaging of ribbon-like condensates formed by co-expressing RIBEYE with Piccolino in HEK293T cells. **d** Domain organization of Piccolino showing multiple PXDLS motifs in its C-terminal region. Multi-sequence alignment of this region indicates that these PXDLS motifs are highly conserved across species. The residue numbers between neighboring PXDLS motifs are indicated in parentheses. The designed constructs of n-PXDLS containing different numbers (n) of PXDLS motifs are illustrated. **e** Regular spatial distribution of PXDLS-binding sites on one face of the B’ sheet. Distances between neighboring PXDLS-binding sites on adjacent B’ filaments are indicated by dashed lines, with blue lines connecting proximal sites and red lines connecting distal sites. The distances were measured using residue F602 as a reference point on the PXDLS-binding site of B domains. **f** A cartoon model illustrating the effect of Piccolino in promoting B’ filament formation and bundling. **g** Sedimentation analysis of B’ filament formation with or without n-PXDLS fragments. To maintain consistent concentrations of individual PXDLS motifs in all conditions, fragment concentrations were calculated by dividing 20 μM by the number of PXDLS motifs per fragment (n). The boundaries used for n-PXDLS fragments are illustrated in panel **d**. **h** Quantitative analysis of B’ filament formation in the presence of various concentrations of n-PXDLS fragments. Repeated 3 times for each group. The estimate of variation is indicated by the s.d.. The corresponding raw SDS-PAGE data are shown in Supplementary Fig. S10d. **i** NsEM analysis of the promoted B’ filament formation and bundling by the addition of 6-PXDLS. For the sample preparation, 20 μM B’ was mixed with 6-PXDLS at a molar ratio ([B’]/[PXDLS]) of 1:1. Before the NsEM analysis, the sample was diluted to ∼1 μM. **j** Cell imaging of condensates formed by co-expressing RIBEYE with Piccolino truncations in HEK293T cells. The first 4 or 6 PXDLS motifs were removed from Piccolino to generate the truncation constructs. **k** Quantification of condensate morphology in cells shown in panel **j**. n = 3 repeats, in each repeat 110-130 cells are quantified. The estimate of variation is indicated by the s.d..

To determine the regulatory mechanism of Piccolino in the morphological transition of RIBEYE condensates, we co-expressed RIBEYE with Piccolino in cells. Consistent with previous observations ^40^, Piccolino transformed the spherical shape of RIBEYE condensates into ribbon-like structures (Fig. 5c). This transformation is likely mediated by the C-terminal region of Piccolino, which contains multiple PXDLS motifs (Fig. 5d; Supplementary Fig. S10b). The binding of these PXDLS motifs to the individual B domain may stabilize neighboring B domains to facilitate the high-order assembly of B’ filaments (Fig. 3e; Supplementary Fig. S10c). In our RIBEYE_B’ sheet model, the neighboring PXDLS-binding sites show a regular spaced arrangement, with inter-filament pairs separated by alternating distances: a short distance of ∼47 Å and a longer distance of 58-71 Å (Fig. 5e). Interestingly, the PXDLS motifs of Piccolino are also arranged in a periodic pattern, with alternating short (∼25 residues) and long (∼45 residues) unstructured spacers (Fig. 5d). Such motif arrangement may bridge two adjacent B’ filaments and promote inter-filament bundling (Fig. 5e). Thus, Piccolino may promote the ribbon-like architecture of RIBEYE through stabilizing the B’ sheet via its periodic spaced PXDLS motifs (Fig. 5f).

To verify our hypothesis, we added purified Piccolino fragments containing varying numbers of PXDLS motifs (designated as n-PXDLS, where *n* denotes the motif count, Fig. 5d) to RIBEYE_B’ and used ultracentrifugation to separate B’ assemblies by molecular size. To maintain a consistent stoichiometric ratio between PXDLS motifs and B’ across different Piccolino fragments, we applied PXDLS motif concentration ([PXDLS], rather than fragment concentration; for example, the molar concentration of 6-PXDLS was 1/6 of that of 1-PXDLS) in the ultracentrifugation analysis. Our result showed that the B’ fragment alone predominantly remains in the soluble fraction, presumably due to its short filament formation in solution. In contrast, the addition of 6-PXDLS dramatically increased the pellet proportion of B’ (Fig. 5g), supporting the effect of 6-PXDLS on the high-order assembly of B’. As the number of PXDLS motifs in the fragments decreases, the ability to promote B’ assembly declines substantially, and the 1-PXDLS motif completely loses its capacity to promote B’ assembly (Fig. 5g, h; Supplementary Fig. S10d). Our nsEM analysis further showed robust B’ filament formation in the presence of 6-PXDLS (Fig. 5i). In contrast to short and curved filaments formed by B’ alone, the 6-PXDLS-promoted B’ filaments are much longer and can even bundle together (Fig. 5i), confirming the strong effect of 6-PXDLS in crosslinking B’ filaments. Consistent with our structural and biochemical findings, our cellular analysis showed that the deletion of six PXDLS motifs from Piccolino severely impaired its capacity to promote the formation of ribbon-like condensates of RIBEYE (Fig. 5j, k).

### CtBP1 promotes RIBEYE phase separation and synaptic ribbon deformation by weakening the B’ filament assembly

CtBP1, another CtBP family member originally found to be in the nucleus, has also been detected in ribbon synapses, where it colocalizes with RIBEYE (Supplementary Fig. S11a) ^48^. Although CtBP1 has a long splicing variant that contains a slightly extended sequence at the N-terminal region, it lacks an RRT motif found in the linker of RIBEYE (Fig. 6a), suggesting that CtBP1 is unlikely to form filaments on its own. Our immunofluorescence staining analysis of the mouse retina confirmed that CtBP1 is exclusively localized on synaptic ribbons in the retina sections (Fig. 6b). Interestingly, CtBP1 levels in the mouse retina increase significantly in mice exposed to light conditions compared to those in the dark (Fig. 6b, c). Given previous observations that light exposure shifts synaptic ribbon morphology from ribbon/horseshoe-shape to droplet-like structures ^26–28,49^, this reciprocal correlation suggests that the CtBP1 level modulates the structural plasticity of synaptic ribbons in response to varying light environments.

**Figure 6.**
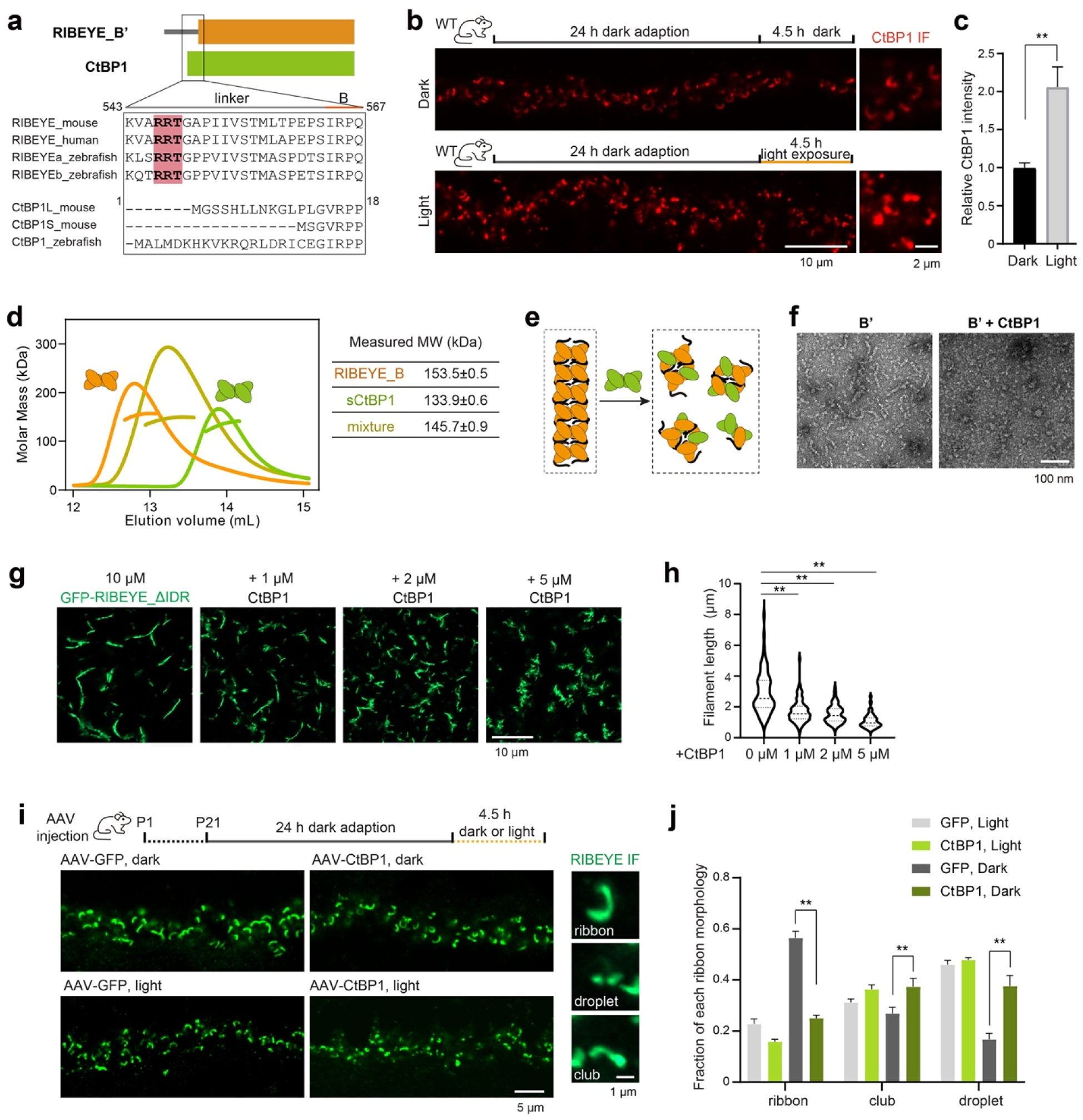
CtBP1 promotes liquid-like condensate formation by disrupting B’ filament formation. **a** Amino acid sequence comparison of RIBEYE_B’ and CtBP1. The highlighted RRT motif in the linker region is unique to RIBEYE. **b** Representative images of the retinal OPL from light or dark-adapted mice. Sections were stained with the CtBP1 antibody. **c** Quantification of the intensities of CtBP1 staining in panel **b**. The intensities were normalized by RIBEYE staining. n = 4 animals. The estimate of variation is indicated by the s.e.m.. Student’s t test. **p < 0.01. **d** MALS assay showing the heterotetramer formation of RIBEYE_B and sCtBP1. The theoretical MWs of RIBEYE_B and sCtBP1 are 46.6 kDa and 38.5 kDa, respectively. **e** NsEM analysis of B’ filament formation in the addition of CtBP1 at a 1:1 molecular ratio. **f** A cartoon model illustrating the disruptive effect of CtBP1 on B’ filament formation by forming heterotetramers. **g** Representative images of the ribbon-like structure formed by GFP-RIBEYE_ΔIDR co-incubated with CtBP1. **h** Quantification of the length distribution of the ribbon-like structure in panel **g**. Dotted lines on the violin plot show the medians and the first/third quartiles for each group. n = 70-110 ribbon-like structures. ANOVA followed by Dunnett’s multiple comparisons. **p < 0.01. **i** Representative images of the retinal OPL from light or dark-adapted mice with EGFP or CtBP1 overexpression. Sections were stained with RIBEYE antibody. Zoomed-in views on the right panel show the criteria for quantification of the ribbon morphology. **j** Quantification of the ribbon morphology in panel **i**. n = 4 animals. The estimate of variation is indicated by the s.e.m.. ANOVA followed by Tukey’s multiple comparisons. *p < 0.05, **p < 0.01.

Considering the high sequence similarity (>90%) between the tetramerization interfaces at the CtBP domains of CtBP1 and the B domains of RIBEYE, we propose that CtBP1 may form a heterotetramer with RIBEYE and interfere with its high-order assembly. Guided by prior structural studies of CtBP1 ^50^, we purified a short version of CtBP1 (sCtBP1) that lacks its unstructured regions at both termini. This allowed us to distinguish between the RIBEYE_B tetramer (theoretical molecular weight of 186 kDa) and the sCtBP1 tetramer (154 kDa) using size exclusion columns. Through analytical size exclusion chromatography coupled with multiangle light scattering (SEC-MALS), we determined that both RIBEYE_B and sCtBP1 form stable homotetramers (Fig. 6d). When RIBEYE_B and sCtBP1 were mixed, the two proteins formed complexes with a measured molecular weight intermediate between those of the B tetramer and the sCtBP1 tetramer (Fig. 6d), indicating that CtBP1 can interact with RIBEYE to form a B/CtBP1 heterotetramer. Since CtBP1 lacks the RRT motif essential for B’ filament assembly, this hetero-tetramerization is expected to block the B’ filament formation (Fig. 6e). Indeed, when incubated with CtBP1, the B’ fragment failed to form filamentous structures, as revealed by nsEM analysis (Fig. 6f). Additionally, the length of ribbon-like structures formed by purified GFP-RIBEYE_ΔIDR was significantly decreased by the addition of CtBP1 in a dose-dependent manner (Fig. 6g, h), confirming that CtBP1 perturbs the high-order assembly of RIBEYE.

To test whether CtBP1 plays a role in regulating the deformation of synaptic ribbons *in vivo*, we employed adeno-associated virus (AAV)-mediated retinal injection in mice (Supplementary Fig. S11b). This approach was used to overexpress CtBP1 in photoreceptor cells, mimicking the elevated CtBP1 levels observed following light stimulation. After the virus injection, the majority of the CtBP1 protein accumulated in the outer plexiform layer (OPL), suggesting that CtBP1 tends to localize to synapses (Fig. 6i; Supplementary Fig. S11b). Consistent with our biochemical findings, the overexpression of CtBP1 in the dark-adapted groups caused a significant reduction in the number of ribbon structures while leading to an increase in spherical droplets (Fig. 6i, j). This morphological alteration phenocopies the change observed after light stimulation (Fig. 6b). Thus, our results indicate that CtBP1 is involved in the retinal transition of synaptic ribbons to synaptic spheres by preventing B’ filament extension. Notably, previous studies have shown that genetic ablation of CtBP1 exerts minimal effects on the formation and function of retinal ribbon synapses, although CtBP2 may compensate for the loss of CtBP1 ^51^. Therefore, CtBP1 may not be the sole regulator for ribbon morphological remodeling under dark-light transitions.

## Discussion

The plasticity in morphology and function of membraneless apparatuses is pivotal for cells to adapt to dynamic environmental cues. Our study unveils a molecular logic underlying the dynamic formation and remodeling of synaptic ribbons, a process critical for sensory synapses to tune neurotransmission efficiency in response to dynamic signal inputs such as light and sound. At the core of this adaptability is RIBEYE, a single protein capable of orchestrating two distinct mesoscale architectures, rigid ribbon-like assemblies and fluid droplet-like condensates, through dual actions of phase transition and phase separation. Both types of RIBEYE condensates are likely assembled from a shared nanoscale building block: filamentous clusters formed by the SAM and B’ domains (Fig. 4h). During retinal synapse development, nascent RIBEYE nanoclusters serve as the initial assembly basic units (Fig. 7, left panels). These clusters can undergo phase transition into ribbon structures via directional polymerization and crosslinking, processes enhanced by the synaptic protein Piccolino, which stabilizes filamentous networks, particularly in dark environments where sustained neurotransmitter release is essential for dim-light vision (Fig. 7, middle panels). On the other hand, they may also undergo phase separation into droplet-like condensates through nondirectional interactions mediated by RIBEYE’s IDR. In response to varying light conditions, the presynaptic level of CtBP1 acts as a molecular switch, where increased CtBP1 levels weaken RIBEYE filament integrity, driving the equilibrium toward droplet-like condensates and reducing photoreceptor sensitivity under bright light conditions (Fig. 7, right panels). It is noteworthy that, given the diverse environmental cues that drive its bidirectional transitions, the morphological transition of synaptic ribbons may require additional regulators beyond Piccolino and CtBP1. Critically, the dual ability of RIBEYE to undergo both phase separation and phase transition enables synaptic ribbons to achieve an adaptive functional balance between solid ribbon architectures and liquid-like spherical condensates. This plasticity is fundamentally distinct from pathological protein aggregation, where irreversible and uncontrolled phase transition drives diseases. For example, in neurodegenerative disorders, proteins like α-synuclein and FUS undergo aberrant, irreversible liquid-to-solid phase transitions, forming toxic fibrillar amyloid aggregates that disrupt neuronal functions ^52,53^.

**Figure 7.**
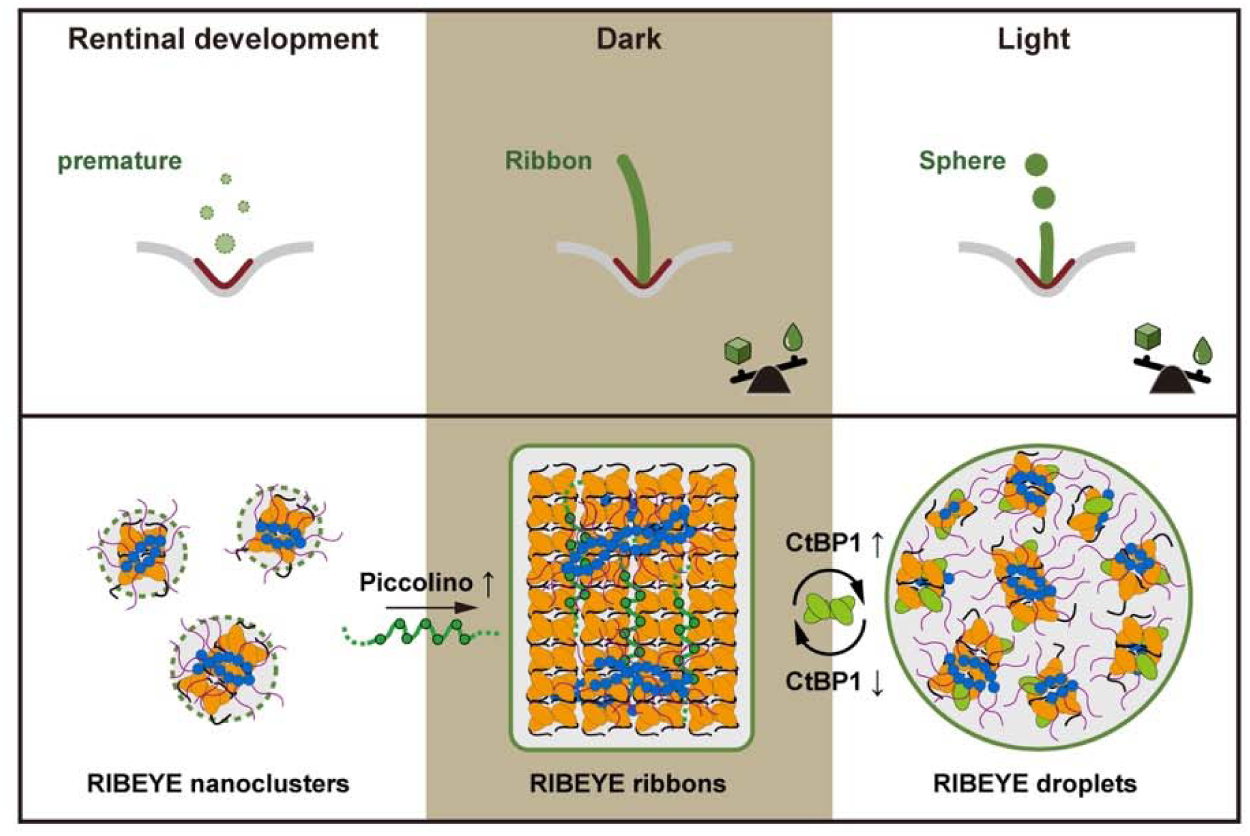
A schematic model illustrating the RIBEYE-mediated phase transition and separation in the plastic assembly and remodeling of retinal synaptic ribbons. RIBEYE molecules and their nanoclusters are depicted as in Fig. 4i. The PXDLS motifs in Piccolino are highlighted as solid circles. These motifs simultaneously bind to multiple B domains, mediating inter-filament crosslinking to drive RIBEYE ribbon assembly from nanoclusters. CtBP1 forms heterotetramers with the B domain of RIBEYE, disrupting B’ filament elongation and promoting RIBEYE droplet-like condensate formation. The required balance between solid and liquid phases of RIBEYE assemblies under different light conditions is schematically represented by seesaws.

Beyond retinal synapses, our findings suggest a generalizable framework for how membraneless apparatuses achieve structural and functional plasticity: nanoscale building blocks (e.g., RIBEYE oligomers) form and dynamically switch between ordered mesoscale architectures and less ordered condensates. This adaptive logic may extend to other cellular contexts, such as cell migration and muscle mechanosensing, where environmental cues like mechanical force drive mesoscale remodeling. For example, focal adhesions (FAs) emerge from nascent adhesion nanoclusters of integrins to form mature, micron-scale adhesions to the extracellular matrix ^54,55^. Their size and shape are sensitive to mechanical forces: low-force environments promote small, dynamic adhesions with liquid-like properties, while high-force conditions stabilize large, rigid adhesions. Recent studies reveal that key FA proteins, including FAK, talin, and paxillin, undergo LLPS to form condensates ^19,20,56^. During FA maturation under high force, actin filaments and FA proteins become highly ordered ^57^, and molecular turnover slows dramatically ^58^, suggesting a force-induced phase transition from liquid-like condensates to solid-like filamentous networks. This plastic assembly mechanism likely governs another mesoscale membraneless apparatus, the Z-disc of muscle sarcomeres, which is critical for actin filament anchoring and force transmission ^16^. During myofibril formation, Z-bodies, nanoscale complexes of α-actinin, titin, and FATZ, grow and fuse to form mature Z-discs ^59,60^. Interestingly, FATZ forms condensates via phase separation and is able to co-condense with α-actinin ^21^, presumably organizing actin filaments and facilitating the transition from nanoscale Z-bodies to mesoscale Z-discs.

Retinal synaptic ribbons typically exhibit a characteristic curved morphology (Fig. 1c) ^5^, in contrast to the relatively flat nature of the sheet structure found in the ribbon-like architecture (Fig. 3b). This disparity suggests that the ribbon structure formed by RIBEYE is capable of undergoing plastic changes *in vivo*. Our *in vitro* findings already offer insights into the plasticity of these ribbon structures. B’ filaments alone show an inherent structural flexibility, indicated by their short and curved filamentous forms (Fig. 2e, f), likely due to the limited interaction between neighboring B tetramers within the B’ filament structure. In addition, the SAM filament can undergo conformational changes between its two forms (Movie S2). Given that the SAM and B’ filaments align in a near-perpendicular orientation, such changes likely alter the spatial arrangement of B’ filaments. Thus, these dynamic transitions of SAM filaments, in conjunction with the flexibility of B’ filaments, may modulate synaptic ribbons to have a curved shape.

Morphologically, synaptic ribbons have diverse shapes, ranging from the thin ribbons (30-40 nm in thickness) found in retinal photoreceptors and pinealocytes to the thick ribbons (∼55-200 nm) in inner hair cells ^61,62^. This morphological variation suggests that the assembly mode of RIBEYE is regulated differently in various sensory neurons. In our cryo-ET analysis of ribbon-like condensates, B’ sheets can stack to form a multi-layered architecture (Fig. 3a), separated by ∼40 nm gaps (Supplementary Fig. S6a), a distance suited for accommodating the ∼280-residue IDR of RIBEYE. This architectural feature allows the parallel stacking of not only two B’ sheets in retinal synaptic ribbons, but multiple sheet structures to generate thick ribbons in cochlear hair cells. In these ribbon architectures, in addition to the SAM filament, IDR may mediate weak, nondirectional interactions between adjacent sheets, thus acting as inter-sheet “glue” as well as spacers. Another question about synaptic ribbon formation is how the synaptic ribbon can extend from the AZ surface. A plausible mechanism involves Bassoon, a scaffold protein critical for anchoring synaptic ribbons to the presynaptic terminal ^63^. Given that Bassoon itself contains PXDLS motifs known to bind RIBEYE’s B domain ^64^, it is tempting to hypothesize that the Bassoon-mediated interaction with RIBEYE organizes B’ filaments of RIBEYE nanoclusters, nucleating synaptic ribbon formation at and simultaneously tethering the ribbon to the presynaptic membrane. Elucidating the molecular mechanism underlying the PXDLS-binding modes of Piccolino and Bassoon to RIBEYE at a mesoscale level will extend the understanding of the development and function of ribbon synapses.

## Methods

### Plasmids

The mouse *RIBEYE* gene and its truncations were cloned from a cDNA library. The SAM (residues 280-375), linker (residues 436-565), B’ (residues 545-988), and B (residues 577-988) fragments were cloned into the pET SUMO vector, with an N-terminal His_6_-SUMO tag. The ΔIDR, ΔSAM, Δlinker, and ΔB constructs were cloned into the pCAG vector, with a Flag-MBP-GFP tag or a mCherry tag added at the N-terminus. For RIBEYE_IDR purification, an 8-residue sequence (residues 85-92) was deleted from the IDR to prevent cleavage by bacterial proteases. All mutations and truncations were generated by site-directed mutagenesis and confirmed by DNA sequencing. The human CtBP1 gene and its truncation (sCtBP1 fragment, residue 4-354) were cloned into a modified pET32 vector with an N-terminal His_6_ tag for affinity purification. The human Piccolino gene and its truncations were cloned into a pTGFP vector. For cellular assays, constructs with varying numbers of PXDLS motifs removed, including Δ1-4 PXDLS motifs and Δ1-6 PXDLS motifs, were generated. For sedimentation assays, Piccolino C-terminal fragments (n-PXDLS, n=1, 2, 3, 6) containing different numbers of PXDLS motifs were introduced into the modified pET32 vector, with an MBP tag at the N-terminus and a Trx tag at the C-terminus. All truncations were generated via homologous recombination and confirmed by DNA sequencing.

### Cell cultures

HEK293T and HeLa cells were cultured in Dulbecco’s modified Eagle’s medium (DMEM) supplemented with 10% FBS, 1% NEAA, and 1% penicillin/streptomycin (P/S) at 37°C. Cell transfection was performed according to the product’s protocol (Neofect).

HEK293F cells were cultured in 293 Chemically Defined Medium (Union) and maintained at 37°C in an incubator with 5% CO2. Polyethylenimine Linear (PEI) MW40000 (YEASEN) was used for transfection.

### Cell imaging

Live-cell imaging was performed using a Nikon A1R confocal Microscope with an environmental chamber maintained at 37 °C and 5% CO2. Cells were seeded on glass-bottom plates. Each plate of cells was transfected with 1 μg of plasmids of mCherry-RIBEYE or associated mutation/truncation. The images were taken after the cells were transfected around 24h.

For fixed cell imaging, HEK293T cells were cultured on glass coverslips in 12-well plates. After 36h of transfection, cells were fixed with 4% paraformaldehyde (PFA) for 15 min at 37°C. The fixed cells were visualized and recorded with a 100× objective using a Nikon A1R Confocal Microscope.

### Correlative light electron microscopy (CLEM)

For the CLEM experiment, HeLa cells were cultured on gridded glass-bottom dishes (D35-14-1.5GI, Cellvis) and transfected with mCherry-RIBEYE_ΔIDR. Transfected cells were fixed with 2% paraformaldehyde (PFA, 16005, Sigma) in 0.15 M PBS (pH 7.35) for 30 min at room temperature. Cells containing ribbon-like condensates were identified using differential interference contrast (DIC) mode, and their positions on the grid were documented. Fluorescence imaging was performed using a confocal microscope (Nikon A1R), and areas with positive cells were marked for subsequent EM processing. Then, samples were post-fixed in 2.5% glutaraldehyde (111-30-8, SPI Inc.) in 0.1 M PB buffer (pH 7.3) at 4°C overnight. After washing twice with PB buffer and twice with ddH_2_O, samples were subsequently fixed with 1% OsO_4_ (wt/vol) and 1.5% (wt/vol) potassium ferricyanide aqueous solution at 4°C for 40 mins, followed by washing. Cell samples were then incubated in filtered 1% thiocarbohydrazide aqueous solution (223220, Sigma-Aldrich) at room temperature for 10 min, 1% unbuffered OsO4 aqueous solution at 4°C for 40 mins, and 2% UA aqueous solution at room temperature for 1h, with four 10-min rinses in ddH_2_O between each step. Dehydration was performed using a graded ethanol series (30%, 50%, 70%, 80%, 90%, and 100% × 2, 5 min each at 4°C). Samples were then infiltrated with graded mixtures of acetone and resin (21 ml SPI-PON812, 13 ml DDSA, 11 ml NMA, and 1.5% BDMA) using sequential ratios of 3:1, 1:1, and 1:3 (v/v), followed by pure resin immersion. Finally, samples were embedded in pure resin and polymerized for 12 hours at 45°C and 48 hours at 60°C. Serial ultrathin sections (50 nm thickness) were obtained using an ultramicrotome (UC7, Leica) equipped with an AutoCUTS device (Zhenjiang Lehua Technology Co., Ltd). Electron microscope images were acquired using a Helios Nanolab 600i dual-beam SEM (Thermo Fisher) with automated imaging software (AutoSEE).

### Protein expression and purification

mRIBEYE-ΔIDR was expressed in HEK293F. The plasmid was transfected with PEI when the cell density reached 2.0 × 10^6^ cells/ml. After a 72-hour culture, cells were collected by centrifugation at 1000g for 20 min. For protein purification, the pellets were resuspended in a buffer containing 20 mM Tris-HCl, 200 mM NaCl, 1 mM EDTA, 1% Triton X-100, pH 7.5. The resuspended cells were lysed by ultrasonication. The protein was first purified using anti-FLAG affinity chromatography and eluted with 500μM FLAG peptide (DYKDDDDK). It was then loaded onto a Superdex 6 increase column (GE Healthcare) for further purification. The freshly purified protein was concentrated to the desired concentration for subsequent experiments.

The SAM, linker, B, and B’ fragments of RIBEYE, CtBP1, sCtBP1, and Piccolino_n-PXDLS fragments were expressed in *E. coli* BL21(DE3) cells. Harvested cell pellets were resuspended in buffer (50 mM Tris pH 7.5, 500 mM NaCl, 5 mM imidazole) and lysed via high-pressure homogenization. The proteins were purified using Ni^2+^-NTA affinity chromatography and eluted with buffer supplemented with 300 mM imidazole. The eluted proteins were further purified using a HiLoad Superdex 200 pg column (GE Healthcare) pre-equilibrated in buffer (20 mM Tris-HCl, pH 7.5, 100 mM NaCl, 1 mM EDTA, 1 mM DTT). The freshly purified proteins were prepared for subsequent experiments.

### Protein filament preparation

To form the SAM filament, the fresh purified SAM fragment at a concentration of 40 μM was incubated on ice for 1 min. The salt sensitivity of the SAM filament was tested by incubating SUMO-SAM in buffers containing varying concentrations of NaCl. SUMO protease was added to the mixture, which was then incubated at 4°C overnight to allow for tag cleavage and filament assembly. Following incubation, the sample was centrifuged at 20,000g for 30 min. The supernatant was collected, and the pellet was resuspended in the same volume of buffer. Equal volumes of supernatant and resuspended pellet were subjected to SDS-PAGE analysis to assess the impact of salt concentration on the SAM filament.

To prepare the B’ filament, the B’ fragment at a concentration of 20 μM was treated with SUMO protease, followed by an overnight incubation at 4°C. To test the promotion effect of Piccolino_6-PXDLS on B’ filament formation, 20 μM B’ was mixed with 3.3 μM Piccolino_6-PXDLS. The mixture was then incubated in the presence of SUMO-protease for 30 min at 4°C. To test the disruptive effect of CtBP1 on B’ filament formation, the B’ fragment was mixed with CTBP1 at a molar ratio of 1:1, followed by overnight SUMO cleavage at 4°C.

### nsEM

For negative staining, sample concentrations were diluted to 1μM. 4 μL of the samples were dropped on glow-discharged carbon-coated grids. After staining with 2.5% uranyl acetate, grids were air dried and visualized using an HT7700 (HITACHI) transmission electron microscope.

### Isothermal titration calorimetry (ITC)

To quantitatively analyze protein-protein interaction, all proteins were prepared in an identical reaction buffer containing 20 mM Tris pH 7.5, 100 mM NaCl, and 1 mM EDTA. The protein of the syringe was prepared for titrating into the reaction cell. For different proteins, combined with different protein concentrations. Experiments were carried out at 25°C. The resulting data were analyzed using the MicroCal PEAQ-ITC analysis software, applying a one-site binding model to determine the dissociation constant (*K*_d_).

### Cryo-EM SPA

#### Cryo-EM sample preparation

The SAM and B’ filament samples were freshly prepared, and 4 μL of each sample was loaded onto a glow-discharged grid (QUANTIFOIL Cu, 300 mesh, 1.2/1.3). Notably, for the SAM filament sample, an additional thin and continuous carbon layer was pre-coated on the QUANTIFOIL grid on the Leica EM ACE600, aiming to increase the particle number in the holes. After 5-second incubation, the grid with protein sample was quickly blotted with clean filter papers for several seconds and then immediately plunged into liquid ethane. All the procedures for freezing protein samples were performed on a Vitrobot (FEI) instrument under conditions of 4 °C and 95% humidity, and finally, the prepared grids were stored in liquid nitrogen for screening and data collection.

#### Data acquisition

The prepared grids were carefully transferred into a Titan Krios G3/G3i transmission electron microscope (Thermo Fisher Scientific) with an acceleration voltage of 300 kV, equipped with a Post-GIF Gatan K3 Summit direct electron detector for data acquisition. Finally, 3714 movies for the SAM filament and 1320 movies for the B’ filament were collected by using SerialEM 3.7 ^65^ in the super-resolution mode with a pixel size of 0.415 Å and 0.46 Å, respectively. Each movie containing 32 frames was recorded with an exposure time of 2.0 seconds and a total dose rate of 50 e-/Å^2^ in a pre-set defocus range between -1.5 μm and -2.5 μm.

#### Data processing

Both datasets for SAM and B’ filaments were processed in cryoSPARC v3.3.1 software ^66^, with similar procedures containing motion correction, CTF estimation, particle picking, 2D classification, 3D classification, and Helix Refinement.

For the SAM filament, 3714 movies were aligned to generate micrographs at a pixel size of 0.83 Å with Patch motion correction, followed by defocus estimation in the embedded CTFFIND4 program ^67^. After removing the junk micrographs, 3555 micrographs were selected for particle picking in a filament-tracer mode, resulting in ∼17.5 million initial particles picked out. After three rounds of 2D classification to remove junk particles, in surprise, two different forms of SAM filaments, including a thick form (∼938k particles) and a thin form (∼148k particles), were clearly separated for respective data processing. After further 2D and 3D classifications, ∼530k good particles for thick-filament and ∼57k good particles for thin-filament were selected to generate two initial filament maps in Helix Refinement. Furthermore, based on these two maps, their helical twist and helical rise were further estimated by Symmetry search utility in cryoSPARC as well as their D1 symmetric packing were identified, which were then applied to refine the two final filament maps with overall resolution of 3.37 Å for thick filament and 3.39 Å for thin filament, respectively.

For the B’ filament, similar procedures and programs for data processing were performed in cryoSPARC. Briefly, 1320 movies were motion-corrected to produce the micrographs with a pixel size of 0.92 Å for CTF estimation and junk removal, resulting in 1020 good micrographs. After filament-tracer-based particle picking, ∼7.7 million initial particles were extracted for analysis by 2D and 3D classifications. Especially in the process of particle cleaning, we removed the relatively short and bent filament particles based on 3D and 2D classification results under different box sizes to further enhance particle homogeneity. Finally, we selected the long and straight filaments containing ∼113k particles for initial map generation, followed by the calculation of helical twist and helical rise. With the application of this helical information, we obtained the final map with an overall resolution of 3.41 Å for the B’ filament.

In the above processes, a box size of 384×384 pixels was mainly selected to extract particles for both SAM and B’ filaments, except that a bigger box size of 512×512 pixels was temporarily used for removing the relatively bent particles of the B’ filament. However, to save the calculation source, we binned the particles by indicated folds during 2D and 3D classifications. Moreover, to support model building, all three density maps were further optimized in DeepEMhancer ^68^ for the postprocess.

#### Cryo-EM model building and refinement

To construct the atomic structures of the SAM and B’ filaments, initial models were generated based on the density maps. Specifically, the SAM model predicted by AlphaFold2 ^69^ and the Cryo-EM structure of the CtBP2 tetramer (PDB: 6WKW) were fitted into their respective density maps. The fitting was carried out in a rigid-body mode using UCSF ChimeraX ^70^. For the thick form of the SAM filament, the initial model incorporated 34 SAM molecules, while for the thin form, it included 28 SAM molecules. In the case of the B’ filament, the initial model consisted of 5 CtBP2 tetramers. In addition, the linker region extended from the N-terminus of the B domain was manually built in Coot ^71^, guided by the local density. Subsequently, the initial models were manually adjusted in Coot to optimize their fit to the density maps. Real-space refinement was then performed using PHENIX ^72^. The final models were further validated against their corresponding cryo-EM maps. The statistical information for cryo-EM data collection, processing, model refinement, and validation was summarized in Table S1. The figures representing cryo-EM structural information were prepared by using UCSF ChimeraX.

### Cryo-ET

#### Cell Culture on EM Grids

Quantifoil R2/4 gold EM grids (Au, 200 mesh) were placed at the center of glass-bottom culture dishes, plasma cleaned for 30 s at 25 mA, and immediately incubated in 1 ml of 70% ethanol under ultraviolet light for 10 min. Grids were coated with 1 ml of 0.1 mg/ml poly-D-lysine (Thermo Fisher Scientific) overnight at room temperature, washed six times with PBS, and subsequently incubated overnight in complete DMEM medium at 37°C.

Transfected cells were trypsinized, and ∼1,000 cells were seeded onto each grid. Grids were incubated at 37°C with 5% CO_2_ for 10 min to promote initial cell attachment, after which 1 ml of complete DMEM medium was gently added. Live-cell imaging was performed on the grids the following day.

#### Plunge Freezing

Plunge freezing was performed using a Vitrobot Mark IV (Thermo Fisher Scientific) set to 25°C and 100% relative humidity. Grids were mounted in tweezers, submerged for 10 s in complete DMEM supplemented with 9% dimethyl sulfoxide (DMSO), blotted (blot force: 7, blot time: 10 s), and rapidly plunged into liquid ethane.

#### Cryo-FIB Milling

Cryo-lamellae were prepared on an Aquilos 2 DualBeam FIB/SEM microscope (Thermo Fisher Scientific). Samples were sputter-coated with platinum at 15 mA for 30 s, followed by deposition of a protective organometallic platinum layer using the gas injection system for 20 s, and then a second platinum sputter-coating at 15 mA for 30 s. SEM imaging was performed using an acceleration voltage of 2-5 kV and a beam current of 13-50 pA. FIB milling was performed with a fixed acceleration voltage of 30 kV and progressively decreasing ion beam currents. Rough milling was conducted at 0.3 nA, followed by intermediate milling at 0.1 nA and fine milling at 30 pA. The final lamella thickness was ∼150 nm.

#### Data Acquisition

Cryo-lamella grids were imaged using a Titan Krios G4 transmission electron microscope (Thermo Fisher Scientific) operated at 300 keV. The microscope was equipped with a cold field emission gun, chromatic (Cc) and spherical (Cs) aberration correctors, and a post-column Selectris X energy filter. Tilt series were acquired on a Falcon 4 direct electron detector (Gatan) at a magnification of 42,000× (pixel size of 2.109 Å). Data collection was performed using SerialEM with a dose-symmetric tilting scheme ^73^. Tilt series were typically collected over a ±51° range from the initial stage tilt, using 3° angular increments. The cumulative electron dose per tilt series was 80–100 e /Å2, and the defocus was set to -6 µm.

#### Data Alignment and Reconstruction

Frames from each tilt-series micrograph were aligned using MotionCor2 ^74^. The contrast transfer function (CTF) was estimated with Gctf ^75^. Motion correction, CTF correction, and tilt-series reordering were automated using a custom bash script (https://github.com/scai20/Auto_proc_cryoET_micrographs). Tomograms were reconstructed either in the Etomo graphical user interface in the IMOD software ^76^, using weighted back-projection or the simultaneous iterative reconstruction technique (SIRT), or with AreTomo ^77^, employing the simultaneous algebraic reconstruction technique (SART). Tomograms were subsequently 4× binned prior to downstream analysis.

#### STA

Tomograms were processed in RELION 5.0.4 for STA. Particles were manually picked from the reconstructed tomograms and extracted for further analysis. Initial alignment and 3D classification identified a subset of particles corresponding to a characteristic B’ filamentous class. This selected class was subsequently refined using the subtomogram refinement in RELION.

#### Segmentation

Filaments were segmented manually in IMOD and visualized using IMOD or UCSF ChimeraX.

#### Measuring Distances Between Membranes

Filaments were segmented in IMOD, and the coordinates were extracted. Inter-membrane distances were calculated using a custom Python script (https://github.com/scai20/Intermembrane-space). Histograms were created in Python with Matplotlib ^78^ and Seaborn ^79^.

### Sedimentation assay

The experimental system consisted of 20 μM SUMO-B’ and varying amounts of Piccolino_n-PXDLS proteins. For n-PXDLS (n=1, 2, 3, 6) fragments, instead of using the fragment concentration, we calculated the concentration based on the number (n) of PXDLS motifs in the fragments. This approach allowed us to more accurately assess the impact of the PXDLS motifs on the B’ filament assembly and bundling. The different concentrations of PXDLS/B’ molar ratios (1:9, 1:3, 1:1) were applied to explore the influence of PXDLS concentration on B’ filament. After incubating and SUMO tag cutting at 4 for about 1 h, the samples were spun at 150,0000g at 4 for 30 min. Following centrifugation, the supernatant was promptly isolated by pipette. The pellet was resuspended with the same volume of protein buffer. Take the same volume of supernatant and pellet run SDS-PAGE separately. The intensity of bands of interest was quantified using ImageJ/Fiji software.

### Fluorophore labeling of proteins

The fluorescent dye iFluor 488 NHS Ester was dissolved in DMSO at 10 mg/ml for storage. Th protein was first concentrated to 20 μM and exchanged into the protein labeling buffer (100 mM NaHCO3, pH 8.3, 200 mM NaCl, 1 mM EDTA, and 1 mM DTT) by using a HiTrap desalting column. The molar ratio of the fluorescent dye and the protein was 1:1 for each reaction. Th reaction was performed at room temperature and quenched by adding 200 mM Tris after 1 hour. The labeled protein and the fluorescent dye were separated by a HiTrap desalting column. Fluorescence labeling efficiency was determined using NanoDrop.

### FRAP

For the FRAP experiment using purified RIBEYE_IDR, 2% fluorophore-labeled protein mixed with unlabeled protein was used. 3% PEG 8000 was added to the system to induce the LLPS of RIBEYE_IDR. FRAP experiments were carried out on a Zeiss LSM 880 confocal microscope, 10 min after PEG treatment at room temperature. For FRAP experiments using protein condensate in HEK293T cells, each well in a 12-well plate of HEK293T cells was transfected with 0.5 μg mCherry-RIBEYE plasmids. One day after transfection, live cell imaging was performed at 37 °C in 5% CO2 using LSM 980 confocal microscope.

The central 1.5-3 μm-diameter circular areas in the droplet were photobleached for both FRAP experiments using protein or cells. To quantify the FRAP curve, the intensity at the pre-bleach point was normalized to 100%, and the intensity right after the bleaching was set to 0%. A regression equation was used to get the mobile ratio (Mf) and the time constant (τ) for FRAP analysis.

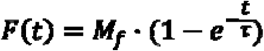

Where F(t) is the percentage of fluorescence recovery and t is the time.

### SEC-MALS

SEC-MALS assays were performed on an AKTA purifier system (Cytiva) coupled with a static light scattering detector (miniDawn, Wyatt) and a differential refractive index detector (Optilab, Wyatt). RIBEYE_B, sCtBP1, and their mixture at a concentration of 100 μM were filtered and loaded into a Superose 6 10/300 GL column pre-equilibrated with a column buffer (50 mM Tris pH 8.2, 100 mM NaCl, 1 mM EDTA, and 2 mM DTT). Data analysis was performed by ASTRA6 (Wyatt).

### Mouse retinal section analysis

#### Mice

Wild-type (C57BL/6J, Charles River) mice of both genders were used. For light or dark adaptation experiments, animals were first adapted to a dark environment for 24 h, followed by room light stimulation or continuous dark adaptation for 4.5 h. Experiments were performed under the guidelines of the Laboratory Animal Facility at the Hong Kong University of Science and Technology.

#### AAV preparation

AAV serotype 2/8 was used for GFP or CtBP1 overexpression in photoreceptor cells. The virus titer was measured by qPCR. The virus titer was 10^13^ GC/mL for all the experiments.

#### Subretinal injection

For surgical procedures involving the neonatal mice, the mice were anesthetized with hypothermia induced by ice embedding. The conjunctiva was cut with the sharp edge of a 30-gauge needle to expose the underlying sclera. 0.5 μL of virus was injected into the subretinal space using a glass micropipette coupled to a Hamilton microsyringe. After surgery, mice were placed on the heating pad to maintain their body temperature until they were fully awake.

#### Immunofluorescence staining

Mice were given a lethal dose of anesthetic and perfused transcardially with PBS, followed by 4% PFA for 5 min for fixation. Mouse retinas were dissected and then post-fixed in 4% PFA for two hours. Retinas were then incubated in 30% sucrose for another two hours and embedded in optimum cutting temperature (OCT) compound. The samples were frozen by dry ice and sectioned at -20 °C. Frozen sections were then permeabilized with 0.1% Triton X-100, blocked with 4% normal goat serum for 30 min, and incubated with respective primary antibodies overnight. After PBS washing three times, sections were incubated with secondary antibodies for two hours and rewashed three times. Primary antibodies, including CtBP2 (BD, 612044), CtBP1 (abcam, ab129181) and Piccolino (abcam, ab20664) were used. Secondary antibodies, including Donkey anti-Mouse IgG AF488 (Invitrogen, A21202), Donkey anti-Mouse IgG AF555 (Invitrogen, A31570), Donkey anti-Rabbit IgG AF488 (Invitrogen, A21206), and Donkey anti-Rabbit IgG AF555 (Invitrogen, A31572), were used.

## Acknowledgments

This work was supported by, National Key R&D Program of China (Grant No. 2025YFA1308800), Fundamental and Interdisciplinary Disciplines Breakthrough Plan of the Ministry of Education of China (JYB2025XDXM505), National Natural Science Foundation of China (82188101 to M.Z, 32471250 to Z.W., 32470576 to S.C., 32400784 to X.W., and 32471262 to F.N.), Key-Area Research and Development Program of Guangdong Province (2023B0303010001), Shenzhen Science and Technology Program (RCJC20210609104333007 to Z.W. and JCYJ20250604144232041 to F.N.), Shenzhen Medical Academy of Research and Translation program (A2303066 to S.C.), Shenzhen Key Laboratory of Biomolecular Assembling and Regulation (ZDSYS20220402111000001), Shenzhen Talent Program (KQTD20210811090115021 to M.Z.), Guangdong Basic and Applied Basic Research Foundation (Grant No. 2024A1515010742 to Z.W., 2023A1515030232 and 2024A1515012651 to F.N.), Guangdong Program (2019QN01Y467 to S.C.), Guangdong Innovative and Entrepreneurial Research Team Program (2021ZT09Y104 to M.Z.), and SUSTech Distinguished Young Scientist Team Program to S.C.. We thank the assistance provided by the Southern University of Science and Technology Cryo-EM Center and Core Research Facilities. Z.W. and S.C. are investigators of SUSTech Institute for Biological Electron Microscopy.

## Author contributions

Z.W. and M.Z. conceived and coordinated the study. Z.W., M.Z., S.C., and K.L. supervised the project. Y.L., XuW., T.Z., and F.N. designed and performed experiments with assistances of C.Y., S.X., Z.Z., W.H., and XuejieW.. Y.L., XuW., T.Z., F.N., Z.S., and K.L. analyzed the data. Z.W., M.Z., and S.C. wrote the manuscript with inputs from other authors.

## Competing Interests Statement

All authors declare that they have no competing interests.

## Data availability

The cryo-EM density maps of the thick SAM filament, the thin SAM filament, and the B’ filament have been deposited into EMDB with accession codes EMD-65841, EMD-65842, and EMD-65843, and their corresponding atomic models have also been deposited in PDB with accession codes 9WBJ, 9WBK, and 9WBL, respectively. The raw cryo-ET tilt series and reconstructed tomograms presented in Figs. 3A and 4H have been deposited in EMPIAR under entry EMPIAR-47487153. The STA density maps of the single- and double-B’ filaments in B’ sheets have been deposited to EMDB with accession code D_9101004548.

---

**Figure S1.**
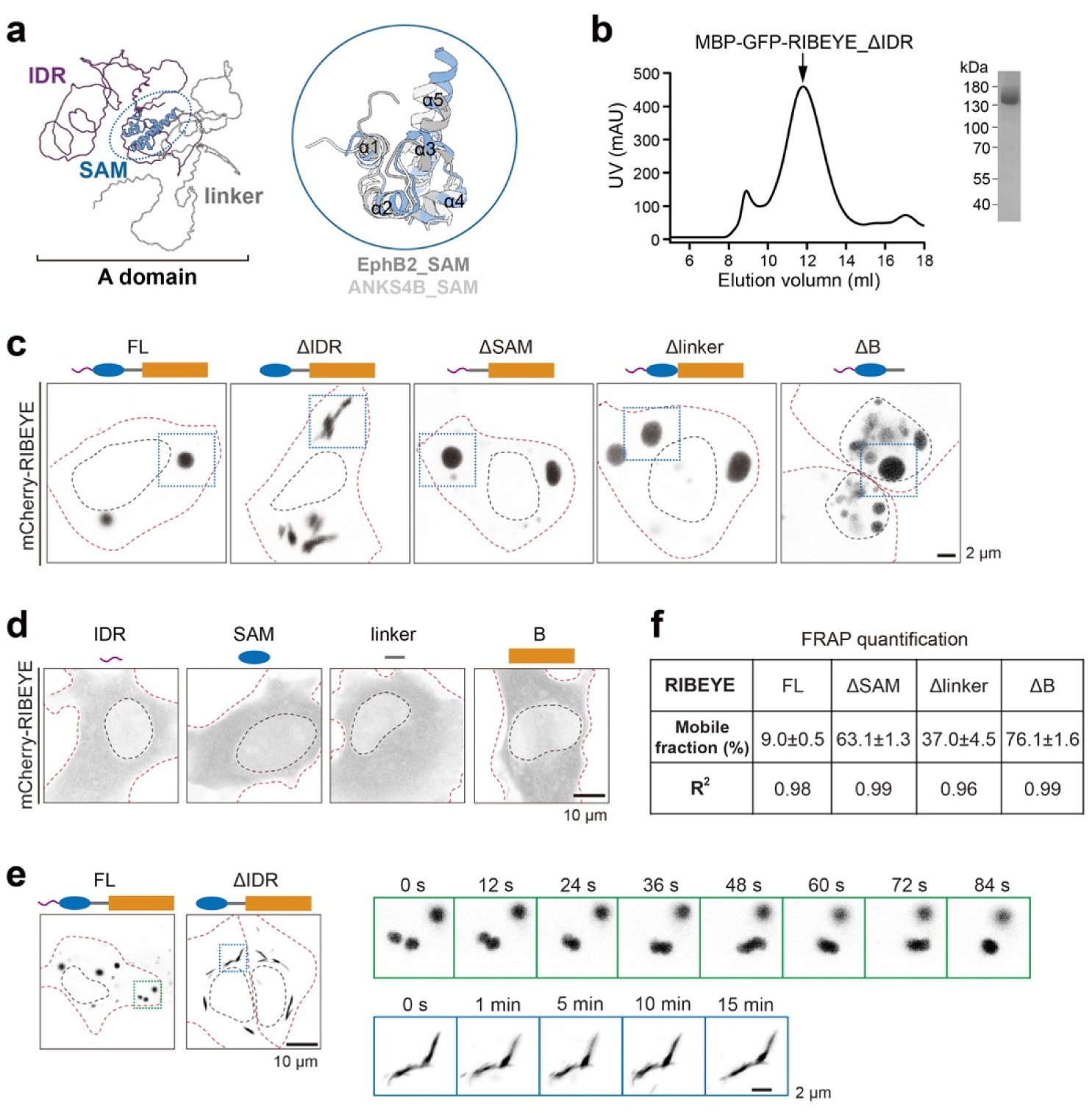
Structural, biochemical, and cellular characterization of RIBEYE. **a** AlphaFold2-predicted structure of the A domain of RIBEYE. The predicted SAM structure of RIBEYE was aligned with the canonical SAM domains in EphB2 and ANKS4B. **b** Size-exclusion chromatographic and SDS-PAGE analyses of the MBP-GFP-tagged RIBEYE_ΔIDR protein. **c** Cell imaging of condensates formed by RIBEYE and its truncations overexpressed in HEK293T cells. The blue dish box indicates condensates used for FRAP analysis, shown in Fig. 1f. **d** Cell imaging of HeLa cells transfected with the IDR, SAM, linker, or B fragments of RIBEYE. **e** Live imaging of spherical and ribbon-like condensates formed by RIBEYE and its ΔIDR truncation overexpressed in HEK293T cells. Different dynamic behaviors of these two condensates were recorded and displayed in the right panels. **f** FRAP quantification analysis of the data presented in Fig. 1g.

**Figure S2.**
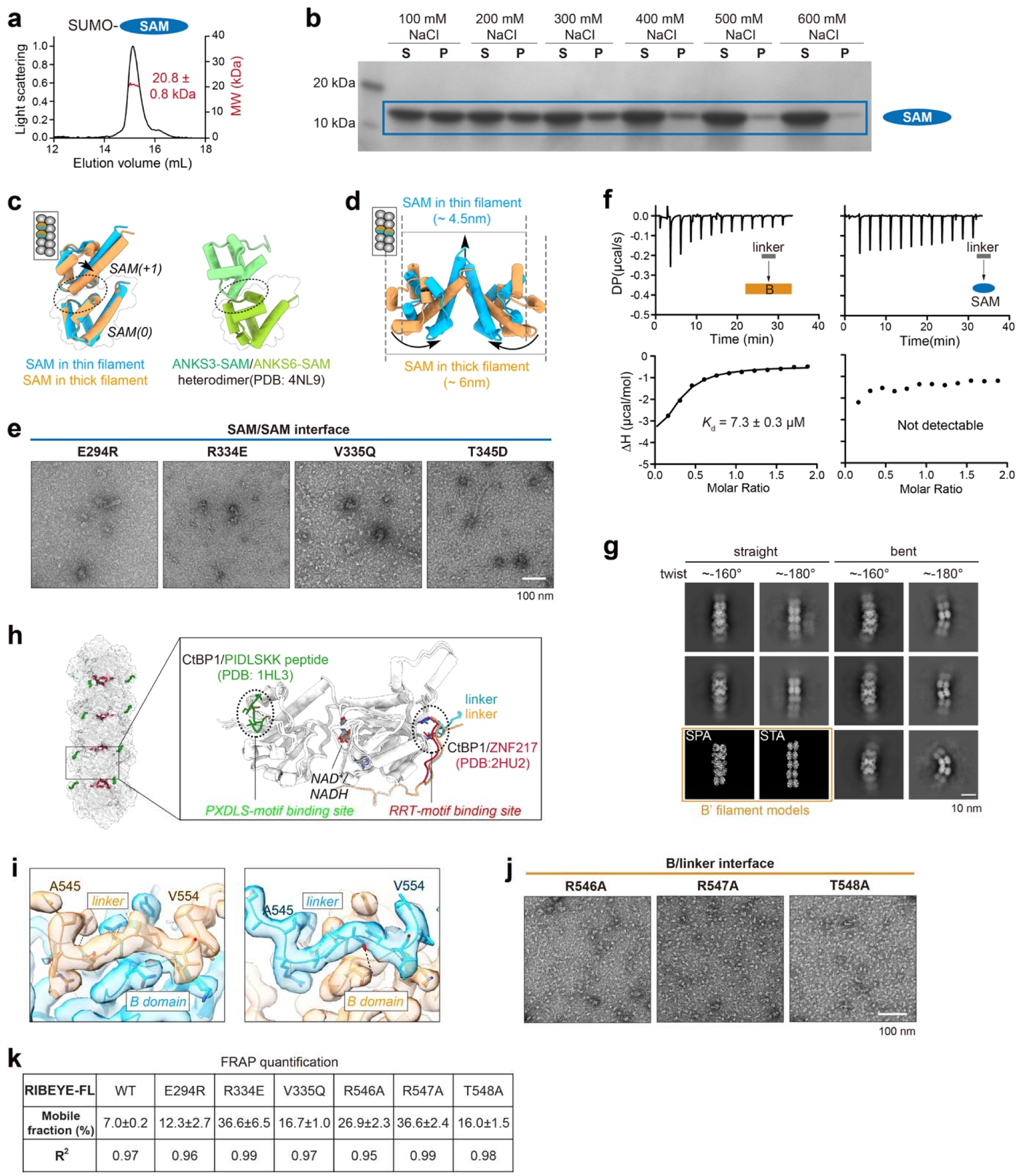
Biochemical and structural analyses of the SAM and B filaments. **a** MALS analysis of the SUMO-tagged SAM fragment (theoretical molecular weight: 22.1 kDa). The protein concentration is 200 μM. **b** Sedimentation analysis of SAM filament formation in the presence of the indicated concentration of NaCl. The SUMO tag was cleaved before sedimentation analysis. With increasing NaCl concentration, the amount of SAM filament in the pellet decreases, indicating disruption of SAM filament assembly. **c** Structural comparison of thick and thin SAM filaments and ANKS3/6-SAM heterodimer. The SAM/SAM interfaces in these SAM structures are circled to indicate a similar interaction mode. **d** Structural comparison of the different inter-helical interactions mediated by α5-helix between the thick and thin forms of the SAM filament. Relative conformational changes between these two forms are indicated by arrows. **e** NsEM analysis showing the disruptive effects of SAM filament formation by SAM/SAM interface mutations. **f** ITC-based analysis of the binding of the linker to the B and SAM domains. The protein concentration of the linker was 400 μM, while the concentration of the B and SAM domains was 40 μM. **g** Cryo-EM analysis of the B’ filament. The 2D classification displayed representative classes of straight and bent filaments with two distinct twist angles. As references, two atomic models of the B’ filament, corresponding to these twist angles, were prepared based on the cryo-EM structure (Fig. 2g) and cryo-ET STA model (Fig. 3c), respectively. **h** PXDLS- and RRT-motif binding sites on the B’ filament, shown by superimposing PIDLSKK peptide (green) and ZNF217 (red) on the filament with alignment of B domains. **i** Cryo-EM densities of the B/linker interface in the B’ filament shown in a model-fit-map mode. The high-quality map allows unambiguous model assignment of the linkers. **j** NsEM analysis showing the disruptive effects of B’ filament formation by B/linker interface mutations. **k** FRAP quantification analysis of the data presented in Fig. 2k.

**Figure S3.**
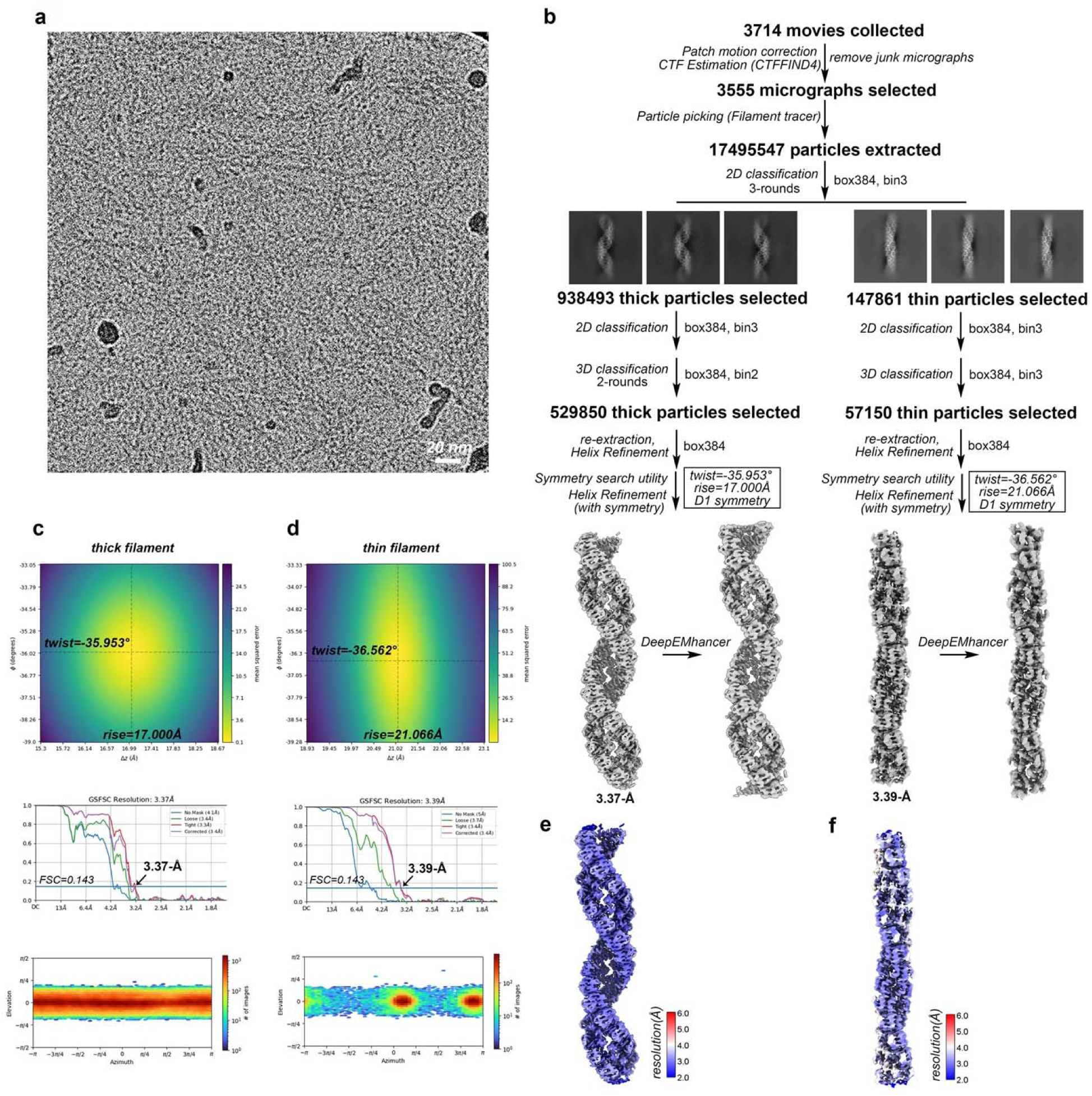
Cryo-EM data processing of the SAM filament. **a** A cryo-EM raw image of the SAM filament. **b** Step-by-step data processing procedures to generate cryo-EM maps of the SAM filament in thick and thin forms. **c-f** The estimation of helical twist and rise, the curves of gold-standard FSC to calculate the map resolution, the distribution of particle orientations, and the map local resolution estimation are represented for thick (**c, e**) and thin (**d, f**) SAM filaments, respectively.

**Figure S4.**
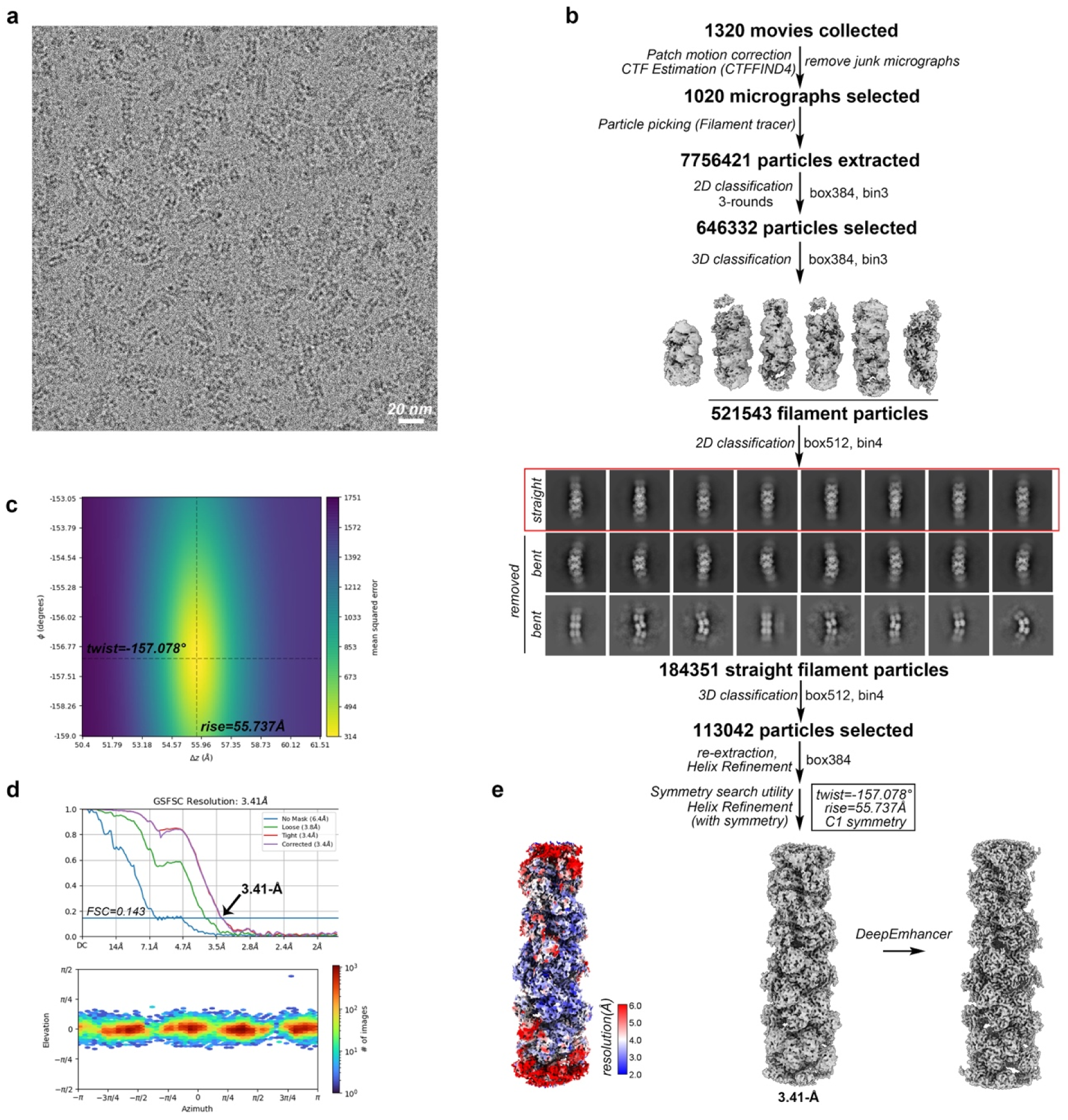
Cryo-EM data processing of the B’ filament. **a** A cryo-EM raw image of the B’ filament. **b** Step-by-step data processing procedures to generate cryo-EM maps of the B’ filament. The relatively straight filament particles boxed in red were selected for the following high-resolution structure determination. **c-e** The estimation of helical twist and rise (**c**), the curves of gold-standard FSC to calculate the map resolution, the distribution of particle orientations (**d**), and the map local resolution estimation (**e**) are represented for the B’ filament.

**Figure S5.**
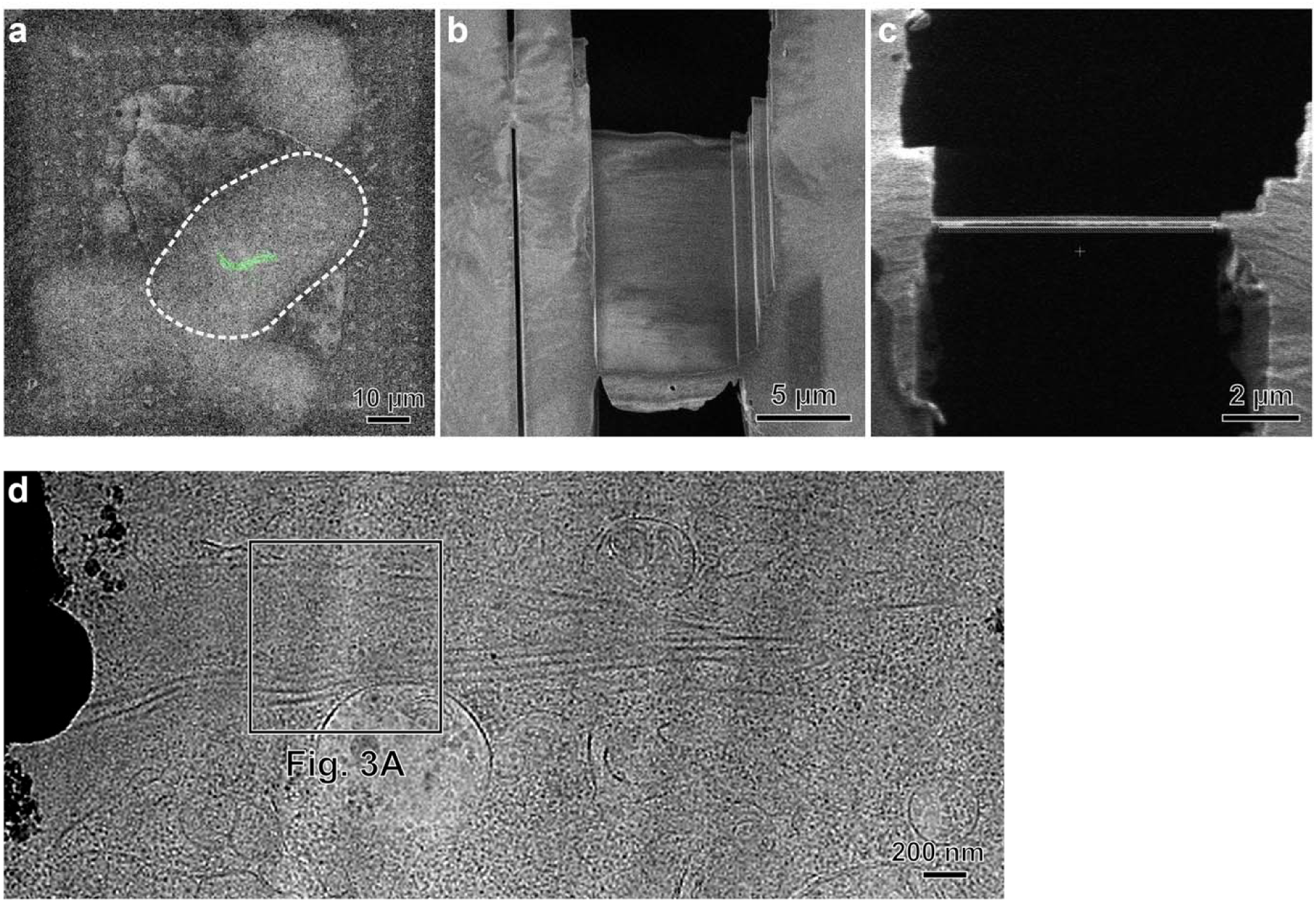
Cryo-ET sample preparation of cells expressing RIBEYE_ΔIDR. **a** SEM image of frozen-hydrated HeLa cells overlaid with a confocal fluorescence image of GFP-tagged RIBEYE_ΔIDR. The GFP signal was used to guide subsequent cryo-FIB milling. The dashed line represents the boundary of a cell. **b, c** SEM view and ion-beam view, respectively, of a ∼150-nm-thick cryo-FIB-milled lamella. **d** Low-magnification cryo-EM projection image of a cryolamella. A cryo-ET tilt series was collected at the boxed region, with the corresponding reconstructed tomogram shown in Fig. 3a.

**Figure S6.**
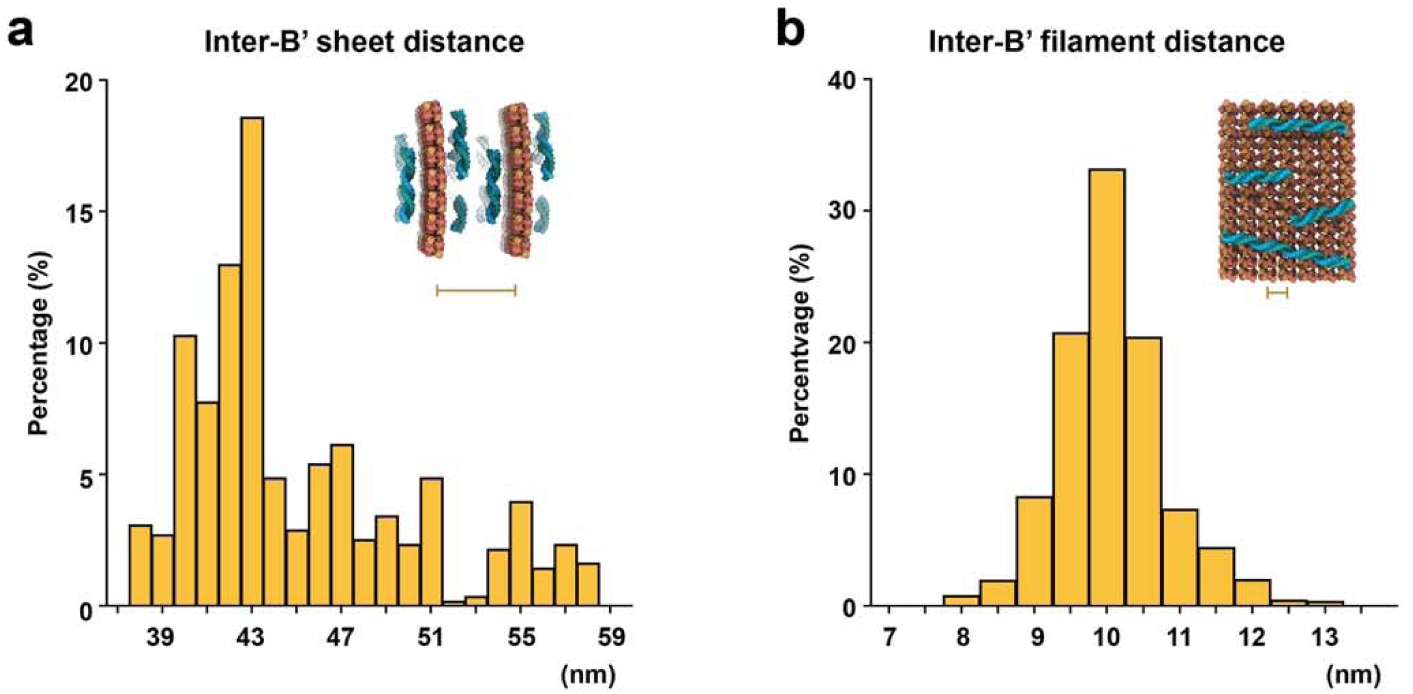
Histogram-based analysis of inter-sheet and inter-filament distances in the ribbon-like condensates formed by RIBEYE_ΔIDR. **a** Quantification of inter-B’ sheet distances. Distances were measured as the center-to-center spacing between adjacent B’ sheets throughout the tomogram. **b** Quantification of inter-B’ filament distances. Distances were measured as the center-to-center spacing between adjacent B’ filaments throughout the tomogram. Data are generated based on cryotomographic slices of GFP-tagged RIBEYE_ΔIDR expressed in HeLa cells.

**Figure S7.**
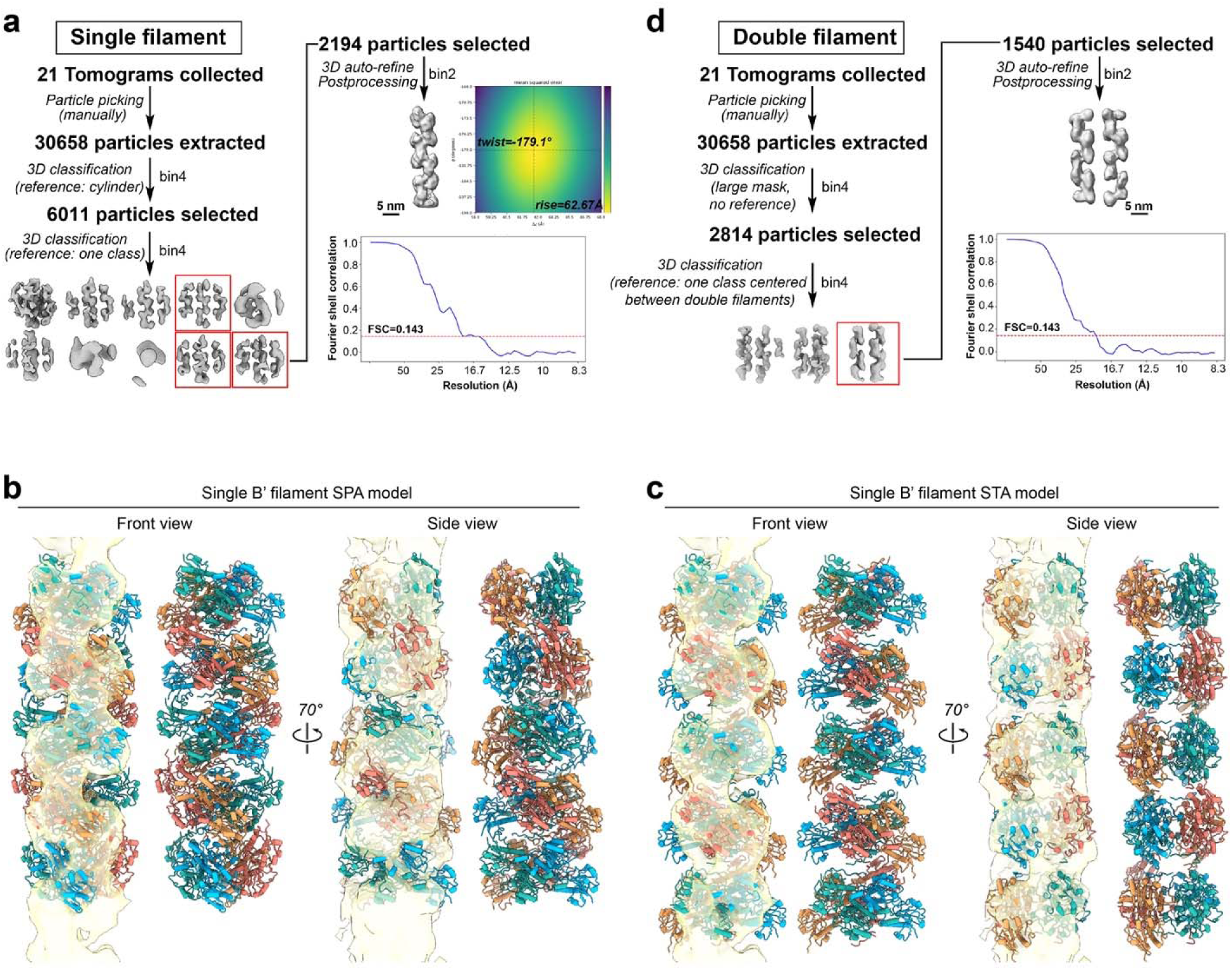
STA workflow and model fitting for single and double B’ filaments. **a** STA workflow for single B’ filaments. The workflow began with a featureless cylindrical reference. A class average or refined map that resembled the single-particle cryo-EM structure of the B′ filament was then selected as the reference for subsequent iterative classification and refinement. Final refinement was performed with binned (2×) data, as unbinned data did not yield further improvements. The helical twist and rise of the STA density were estimated by using Symmetry Search Utility in CryoSPARC. The curve of gold-standard FSC to calculate the map resolution was indicated. **b, c** Superimposition of B’ filament models with the STA density map of the single filament. The single B’ filament models derived from SPA (**b**) and STA (**c**) were each fit into the STA density map of the single filament. For comparison, their top tetramers were aligned in the same orientation within the STA density map. **d** STA workflow for double B’ filaments. No reference was used in the first round of 3D classification. A large mask encompassing three filaments was applied during the initial rounds of 3D classification. Then the densities corresponding to two adjacent B’ filaments from one 3D class were centered within the box and used for further classification and refinement. Final refinement was performed with binned (2×) data, as unbinned data did not yield further improvements. The curve of gold-standard FSC to calculate the map resolution was indicated.

**Figure S8.**
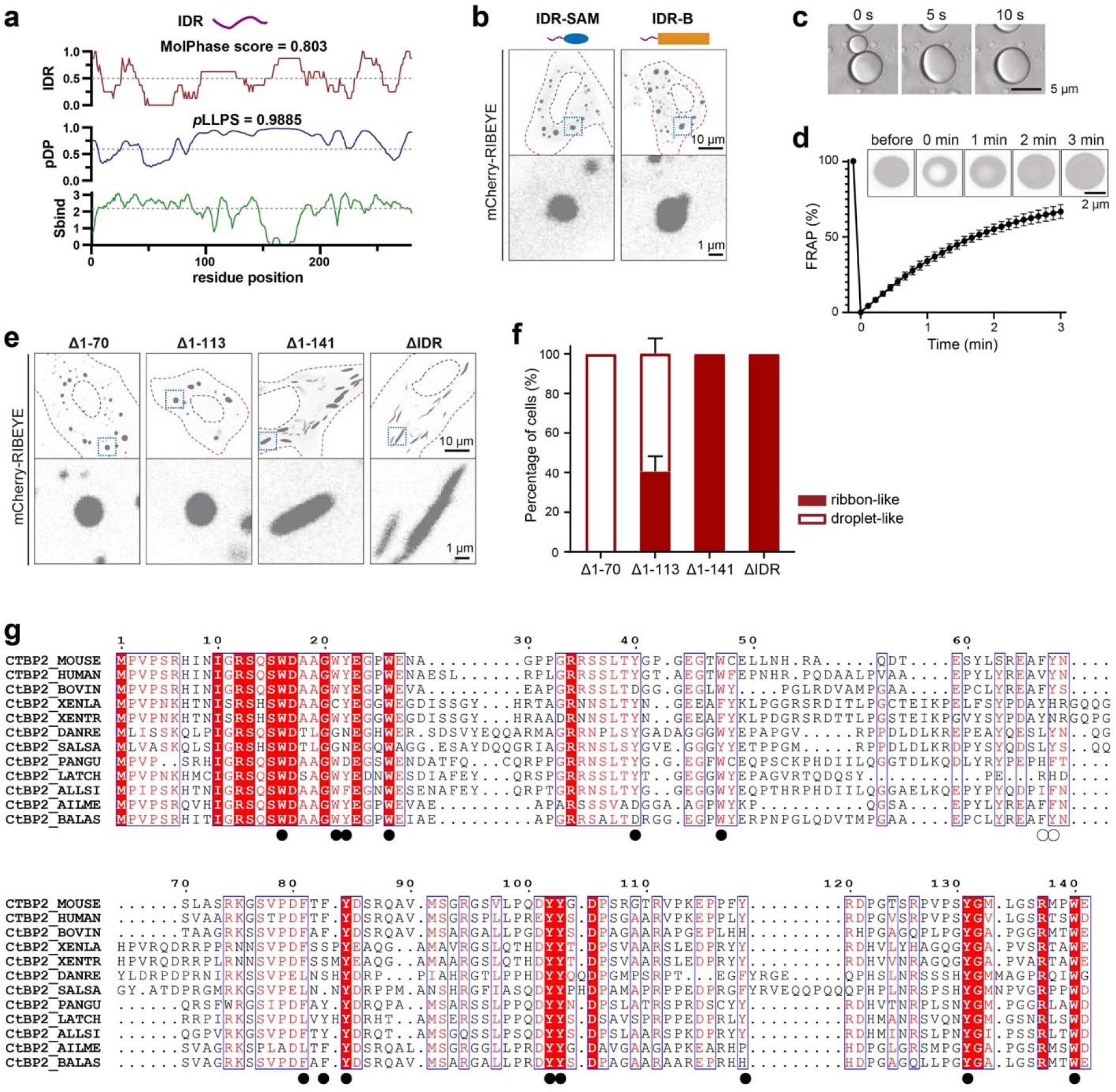
Analyses of the LLPS-promotion capability of the IDR in RIBEYE. **a** Sequence-based prediction of the phase separation probability in RIBEYE_IDR. The residue-based IDR score is calculated using MolPhase (https://molphase.sbs.ntu.edu.sg). The residue-based droplet-promoting probabilities (pDP) and cellular context-dependence (Sbind) are calculated using FuzDrop (https://fuzdrop.bio.unipd.it/). **b** HeLa cell imaging of condensates formed by IDR-SAM and IDR-B, showing that the IDR promotes the transition of both the SAM and B domains from a diffuse cytosolic distribution to droplet-like condensates. **c** Representative images of a fusion event of IDR droplets in the presence of 3% (w/v) PEG 8000. The protein concentration is 40 μM. **d** FRAP analysis of condensates formed by RIBEYE_IDR in the presence of 3% (w/v) PEG 8000. The protein concentration is 40 μM. n = 8 condensates. The estimate of variation is indicated by the s.e.m.. **e** HeLa cell imaging of condensates formed by RIBEYE with different N-terminal truncations. **f** Quantification of condensate morphology in cells shown in panel **e**. n = 3 repeats, in each repeat 20-40 cells are quantified. The estimate of variation is indicated by the s.d.. **g** Alignment of the IDR sequences from different species, including human, bovine, mouse, *Danio rerio*, *Xeenopus laevis*, *Xenopus tropicalis, Latimeria chalumnae, Alligator sinensis, Salmo salar, Balaenoptera acutorostrata, Pantherophis guttatus*, and *Ailuropoda melanoleuca*. The 14 aromatic amino acids mutated in this study are indicated as dark circles.

**Figure S9.**
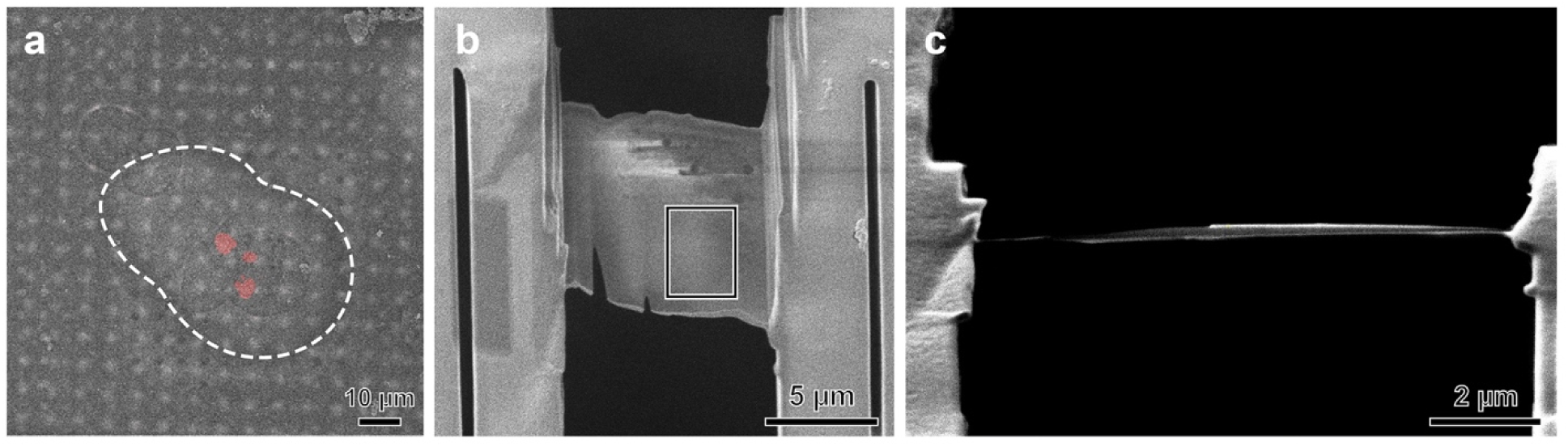
Cryo-ET sample preparation of cells expressing RIBEYE_FL. **a** SEM image of a frozen-hydrated HEK293T cell overlaid with a confocal fluorescence image of mCherry-tagged RIBEYE_FL. The mCherry signal guided subsequent cryo-FIB milling. The dashed line indicates the cell boundary. **b, c** SEM view (**b**) and ion-beam view (**c**) of a ∼150-nm-thick cryo-FIB-milled lamella. The boxed region was selected for high-magnification montage acquisition, with the resulting montage shown in Fig. 4g.

**Figure S10.**
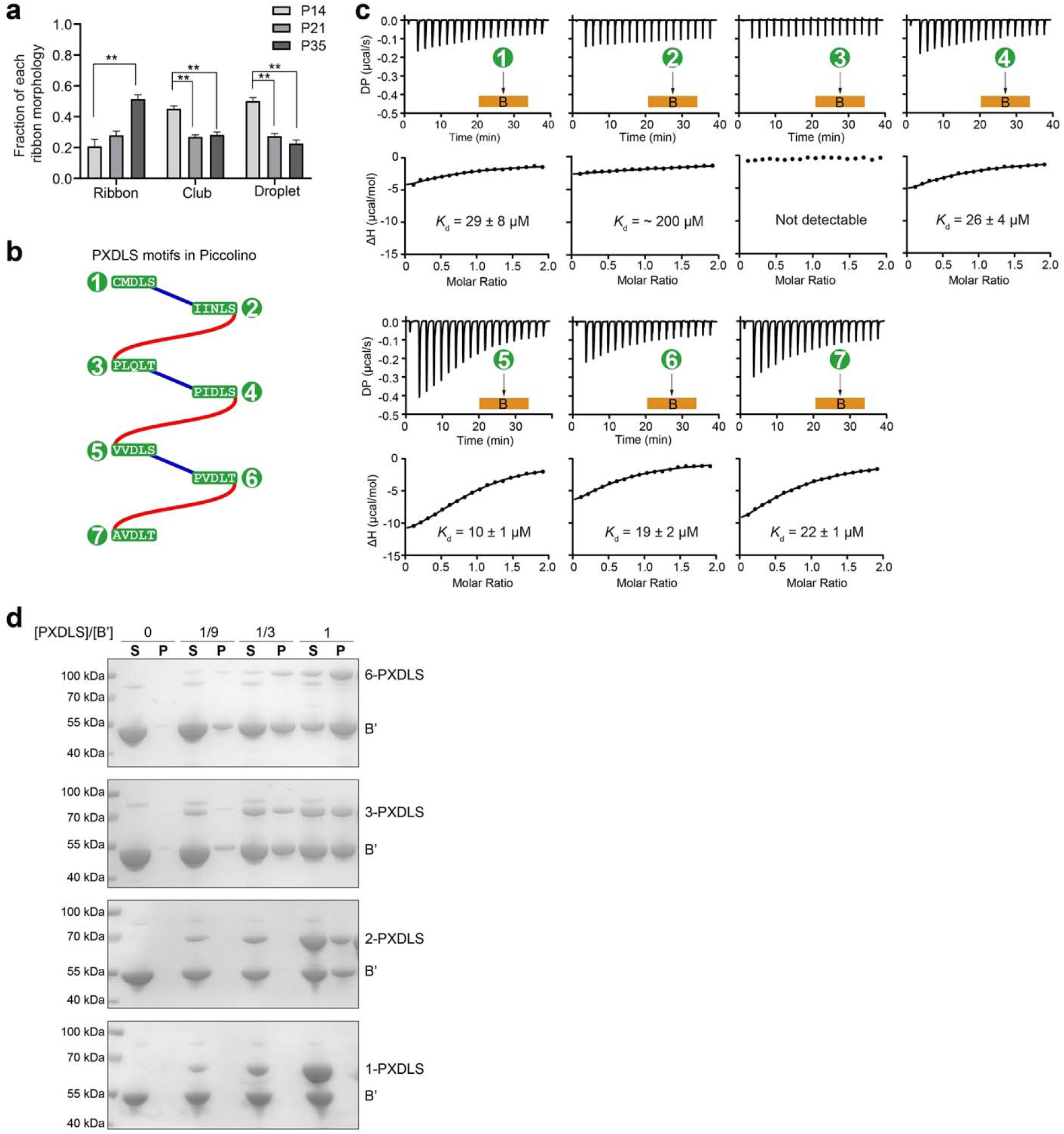
The promotional effect of Piccolino on the RIBEYE-mediated ribbon formation. **a** Quantification of the ribbon morphology shown in Fig. 5a. n = 4 animals. The estimate of variation is indicated by the s.e.m.. ANOVA followed by Tukey’s multiple comparisons. **p < 0.01. **b** Schematic diagram of the seven PXDLS motifs in the C-terminal region of Piccolino. **c** ITC-based analysis of the binding of different PXDLS motifs to the B domain in RIBEYE. **d** Raw SDS-PAGE data for quantification analysis shown in Fig. 5h.

**Figure S11.**
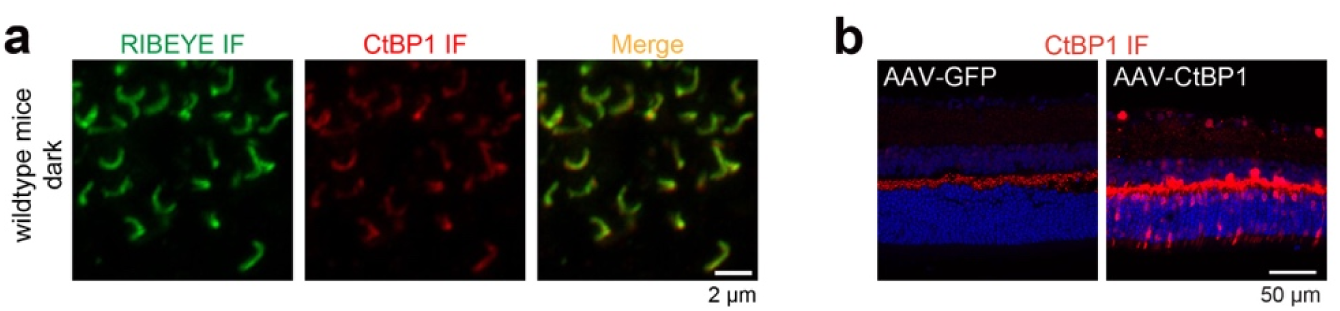
The disruptive effect of CtBP1 on the RIBEYE-mediated ribbon formation. **a** Representative images of the OPL from mice with dark adaptation. Sections were stained with CtBP1 (red) and RIBEYE (green) antibody. **b** Representative images of the OPL from mice injected with AAV-EGFP or AAV-CtBP1. Sections were stained with CtBP1 antibody (red) and DAPI (blue).

**Movie S1. Assembly modes of RIBEYE SAM domains in thick- and thin-filament forms.**

**Movie S2. Conformational transition between thick- and thin-filament forms of RIBEYE SAM domains.**

**Movie S3. Assembly mode of the B’ filament via interactions between B tetramers and N-terminal extended linkers.**

**Movie S4. Cryotomogram and 3D rendering of B’ sheets in a HeLa cell overexpressing RIBEYE_**Δ**IDR.** This subvolume corresponds to the cryotomogram shown in Fig. 3a.

**Movie S5. STA density map and fitted model of a single B’ filament in the ribbon-like condensate of RIBEYE.**

**Movie S6. STA density map and fitted model of double B’ filaments in the ribbon-like condensate of RIBEYE.**

**Movie S7. Cryotomogram and 3D rendering of perpendicularly oriented B’ sheets in a HeLa cell overexpressing RIBEYE_**Δ**IDR.** This subvolume corresponds to the cryotomogram shown in Fig. 3f.

**Movie S8. Structure model of RIBEYE B’ and SAM domains in mesoscale synaptic ribbon assembly.** This model corresponds to the atomic model shown in Fig. 3g.

**Movie S9. Cryotomogram and 3D rendering of B’ sheets in a HEK293T cell overexpressing RIBEYE_FL.** This subvolume corresponds to the cryotomogram shown in Fig. 4h.

**Table S1.** The statistics of cryo-EM data collection, process, model refinement and validation.

| Data collection and process | SAM filament |  | B' filament |
| --- | --- | --- | --- |
| Microscope | Titan Krios G3 |  | Titan Krios G3i |
| Detector | Gatan K3 |  | Gatan K3 |
| Voltage(keV) | 300 |  | 300 |
| Total electron exposure (e- /Å<sup>2</sup>) | 50 |  | 50 |
| Exposure time (s) | 2.0 |  | 2.0 |
| Electron exposure/frame (e- /Å<sup>2</sup>) | 1.56 |  | 1.56 |
| Defocus range (μm) | -1.5 to -2.5 |  | -1.5 to -2.5 |
| Pixel size (Å) | 0.415 |  | 0.46 |
| Initial particle images (no.) | 17495547 |  | 7756421 |
|  | Thick form | Thin form |  |
| Final particle images (no.) | 529850 | 57150 | 113042 |
| Map resolution (Å) | 3.37 | 3.39 | 3.41 |
| FSC threshold | 0.143 | 0.143 | 0.143 |
| Map sharpening B factor (Å<sup>2</sup>) | -158.2 | -128.6 | -51.7 |
| Symmetry imposed | D1 | D1 | C1 |
| Helical twist (°) | -35.953 | -36.562 | -157.078 |
| Helical rise (Å) | 17.000 | 21.066 | 55.737 |
| Refinement |  |  |  |
| Initial model used (PDB code) | AF2-predicted model | AF2-predicted model | 6WKW |
| Model composition |  |  |  |
| Non-hydrogen atoms | 17816 | 14672 | 54720 |
| Protein residues | 2346 | 1932 | 6936 |
| Ligands | 0 | 0 | 20 |
| R.M.S.D. from ideal geometry |  |  |  |
| Bond lengths (Å) | 0.004 | 0.005 | 0.003 |
| Bond angles (°) | 0.709 | 0.792 | 0.562 |
| **Validation** |  |  |  |
| CC_mask | 0.73 | 0.77 | 0.75 |
| Clashscore | 10.61 | 15.56 | 4.61 |
| MolProbity score | 1.54 | 1.70 | 1.59 |
| Poor rotamers (%) | 0 | 0 | 0 |
| Ramachandran plot statistics |  |  |  |
| Favored (%) | 98.51 | 100 | 94.83 |
| Allowed (%) | 1.49 | 0 | 5.17 |
| Disallowed (%) | 0 | 0 | 0 |

## Notes

### Competing Interest Statement

The authors have declared no competing interest.

